# The pentameric complex is not required for vertical transmission of cytomegalovirus in seronegative pregnant rhesus macaques

**DOI:** 10.1101/2023.06.15.545169

**Authors:** Hsuan-Yuan Wang, Husam Taher, Craig N. Kreklywich, Kimberli A. Schmidt, Elizabeth A. Scheef, Richard Barfield, Claire E. Otero, Sarah M. Valencia, Chelsea M. Crooks, Anne Mirza, Kelsey Woods, Nathan Vande Burgt, Timothy F. Kowalik, Peter A. Barry, Scott G. Hansen, Alice F. Tarantal, Cliburn Chan, Daniel N. Streblow, Louis J. Picker, Amitinder Kaur, Klaus Früh, Sallie R. Permar, Daniel Malouli

## Abstract

Congenital cytomegalovirus (cCMV) infection is the leading infectious cause of neonatal neurological impairment but essential virological determinants of transplacental CMV transmission remain unclear. The pentameric complex (PC), composed of five subunits, glycoproteins H (gH), gL, UL128, UL130, and UL131A, is essential for efficient entry into non-fibroblast cells *in vitro*. Based on this role in cell tropism, the PC is considered a possible target for CMV vaccines and immunotherapies to prevent cCMV. To determine the role of the PC in transplacental CMV transmission in a non-human primate model of cCMV, we constructed a PC-deficient rhesus CMV (RhCMV) by deleting the homologues of the HCMV PC subunits UL128 and UL130 and compared congenital transmission to PC-intact RhCMV in CD4+ T cell-depleted or immunocompetent RhCMV-seronegative, pregnant rhesus macaques (RM). Surprisingly, we found that the transplacental transmission rate was similar for PC-intact and PC-deleted RhCMV based on viral genomic DNA detection in amniotic fluid. Moreover, PC-deleted and PC-intact RhCMV acute infection led to similar peak maternal plasma viremia. However, there was less viral shedding in maternal urine and saliva and less viral dissemination in fetal tissues in the PC-deleted group. As expected, dams inoculated with PC-deleted RhCMV demonstrated lower plasma IgG binding to PC-intact RhCMV virions and soluble PC, as well as reduced neutralization of PC-dependent entry of the PC-intact RhCMV isolate UCD52 into epithelial cells. In contrast, binding to gH expressed on the cell surface and neutralization of entry into fibroblasts by the PC-intact RhCMV was higher for dams infected with PC-deleted RhCMV compared to those infected with PC-intact RhCMV. Our data demonstrates that the PC is dispensable for transplacental CMV infection in our non-human primate model.

**One Sentence Summary:** Congenital CMV transmission frequency in seronegative rhesus macaques is not affected by the deletion of the viral pentameric complex.

## Introduction

For expectant mothers and their health care providers, congenital cytomegalovirus (cCMV) infections are a major global concern as they represent the most common infectious cause of birth defects and brain damage including sensorineural hearing loss and neurodevelopmental delays (*1–3*). Worldwide, about 0.7% of newborns are diagnosed with a congenital infection at birth but the incidence rate is correlated to the socioeconomic status of the population with lower to middle income countries indicating a three-fold higher overall case prevalence rate compared to high income countries (*4*). Despite the severe clinical impact and financial burden placed on affected families and the health care system, more than four decades of clinical research have not resulted in an effective, licensed vaccine to prevent cCMV infection. While the reasons for this lack of success are complex and multifaceted, a general gap in our understanding of the virological factors necessary and sufficient for vertical transmission represents a major hurdle for effective cCMV vaccine development (*5, 6*).

While most cCMV transmissions occur in seropositive pregnancies, likely as the result of a re-infection or reactivation event (*1*), overall, a lower congenital transmission rate is observed in seropositive pregnant women compared to age and income-matched seronegative pregnant women (*7*), indicating that pre-existing, CMV specific, adaptive immune responses can provide some level of protection from transplacental transmission. Hence, while initial CMV vaccine trials attempted to induce broad immune responses using serially passaged, live-attenuated HCMV strains (*8*), more recent approaches have largely relied on inducing a strong, neutralizing antibody response against dominant membrane glycoproteins (*3, 4*). The most widely examined HCMV vaccine target is glycoprotein B (gB), a large, highly glycosylated class III viral fusogen found on the viral surface where it mediates fusion of the viral and cellular membrane enabling virus entry (*9*). Recent vaccine approaches also included five individual subunits (UL128/UL130/UL131A/gH/gL) forming the pentameric complex (PC) (*10–13*), one of the two viral entry receptors conserved across all primate CMV species and a target of potent neutralizing antibodies (*14–18*). While the CMV encoded trimer consisting of gH, gL and gO is needed for entry into fibroblasts by cell fusion after binding to platelet-derived growth factor receptor alpha (PDGFRα) (*19, 20*), the PC is needed to facilitate endocytosis into non-fibroblast cells including epithelial, endothelial, and myeloid cells after binding to neuropilin-2 (NRP2) (*21–30*). So far, these cell type specific differences in HCMV receptor usage have solely been observed *in vitro* after infecting primary tissues and cultured cell lines as CMV species-specificity precludes *in vivo* experimentation with the human virus in laboratory animal models. Initial *in vivo* experiments attempted to examine the importance of the PC in RhCMV-seronegative rhesus macaques (RM) by comparing the fibroblast-adapted rhesus cytomegalovirus (RhCMV) strain 68-1 containing multiple mutations and deletions, including subunits of the PC, to the PC-intact, low passage isolates UCD52 and UCD59. Upon s.q. inoculation it was observed that 68-1 displayed restricted tropism at the skin injection site (*31*) and reduced virus shedding in all examined bodily fluids (*32*). However, a more recent study evaluated virus dissemination from the skin injection site after s.q. inoculation of RhCMV-naïve RM with either full-length (FL)-RhCMV, a 68-1-derived, PC-intact clone containing a complete genome, or a FL-RhCMV-derived recombinant that lacked two subunits of the PC and the UL146 family of viral CXC-chemokine like proteins. While significantly reduced spread of the viral deletion mutant was observed throughout the host, the ratio of infected cell types in the spleen was not different (*33, 34*). These mixed results do not allow for a definitive conclusion but could imply that while deletion of the PC clearly leads to virus attenuation *in vivo*, the significant tropism changes observed *in vitro* might not be as pronounced in the infected host. Additionally, novel, PC independent functions for single components have been described, which could contribute to the observed *in vivo* attenuation. UL128 and UL130 are involved in immune programming as each protein can independently inhibit the priming of unconventionally MHC-E and MHC-II restricted CD8+ T cells (*35, 36*), while UL131A has been demonstrated to be non-essential for *in vitro* culture in fibroblasts but indispensable for infection *in vivo* in RM, suggesting a critical role in immune evasion during the initial stages of infection (*37*).

*In vitro*, the PC represents a major target of neutralizing antibodies (NAb) and high level PC-specific humoral immune responses have been described in persistently HCMV infected, healthy individuals, in plasma derived hyperimmune globulin (HIG) solutions, and are potently-induced via immunization with CMV vaccine candidates expressing a complete PC or individual PC components (*14–18*). Yet, the question of whether the complex is required for infection of the placenta and fetal infections remains undetermined. A comprehensive single-cell transcriptome study profiling cell populations in human placenta collected during the first trimester of pregnancy identified various immune and non-immune cell types at the maternal-fetal interface (*38*). As infection of most of these cells are expected to require HCMV to express a functional PC based on *in vitro* studies, it is conceivable that the PC is necessary for transplacental CMV transmission *in vivo*. This hypothesis is supported by the observation that in HCMV seronegative pregnant women a delayed IgG response against the PC and its subcomponent gH/gL within the first 30 days after the primary infection is correlated with intrauterine HCMV transmission (*39*). Additionally, published results using the relevant guinea pig CMV (GPCMV) congenital transmission model demonstrated that restoration of the PC in a lab adapted strain carrying inactivating mutations resulted in increased transplacental transmission and pub mortality directly linking the PC to cCMV infection and fetal pathology (*40*). These observations hint at a critical role for the PC in cCMV transmission and warrant further examinations in an animal model that closely mimics human cCMV pathogenesis.

In this study, we aimed to determine whether the PC is important for CMV transplacental transmission to inform future cCMV vaccine development. Co-evolution of CMVs with their respective host species for millions of years has resulted in strict species-specificity (*41*). Thus, cCMV transmission studies require the use of a relevant animal model recapitulating human pregnancy and immunity in combination with the animal-specific CMV species. Our established cCMV model examines seronegative, pregnant RM inoculated with minimally passaged, pathogenic RhCMV strains. By deleting the PC subunits Rh157.5 (UL128) and Rh157.4 (UL130) from FL-RhCMV, a wildtype-like, bacterial artificial chromosome (BAC)-derived RhCMV clone (*33*), we generated a mutant that specifically lacks the PC. We observed that this virus can still replicate at and disseminate from the skin injection site after s.q. inoculation, albeit to reduced levels compared to the parental control. Intriguingly, viral DNA was detected in the amniotic fluid and fetal tissues in animals infected intravenously (i.v.) with the PC-intact as well as the PC-deleted RhCMV and the overall transmission rates between the two viruses were identical. These results imply that the PC is not required for cCMV transmission in RM, the evolutionary closest, usable primate model to humans. These results are surprising given the demonstrated importance of the PC for cell tropism *in vitro*, especially for the cell types found at the maternal-fetal interface and suggest that the tropism importance observed in tissue culture model systems is not necessarily predictive of viral dissemination and congenital transmission *in vivo*. While our data do not exclude the PC as a useful component in CMV vaccine formulations, they do point to a less significant role in transplacental transmission of PC-specific immunity than originally expected based on *in vitro* and small animal model results and raise the question whether other viral targets more directly involved in cCMV infection need to be identified and interrogated to improve upon existing vaccine approaches.

## Results

### Deletion of the PC results in restricted cell tropism *in vitro* and diminished dissemination *in vivo*

We recently published the construction and characterization of FL-RhCMV (*42*), the first BAC-derived RhCMV clone containing a complete genome. FL-RhCMV was generated by repairing multiple mutations, gene deletions, and inversions that the 68-1 strain had acquired during prolonged tissue culture (*43*). These tissue culture adaptations included the inversion of a gene segment in the genomic locus homologous to the ULb’ region in HCMV. As a result, the two PC subunits Rh157.5 and Rh157.4 and three of six members of the UL146 family of viral CXC chemokine-like proteins were deleted (*35, 44*). Unlike strain 68-1, FL-RhCMV efficiently infected epithelial cells *in vitro*, showed significant replication and shedding in bodily fluids after i.v. inoculation and increased dissemination from the skin infection site to peripheral tissues and organs after s.q. inoculation of CMV-naïve, immunocompetent RM(*33*). The observed peak levels and kinetics of FL-RhCMV genome copy numbers detected in plasma, urine, and saliva after i.v. inoculation was similar to previously used non-clonal, minimally passaged RhCMV strains (*33*), thus indicating that infection by FL-RhCMV is equivalent to wild-type RhCMV strains and can thus be used as a genetically tractable model for RhCMV infections (*42*).

To study the role of the PC in primary and congenital infection we deleted two of the five subunits, open reading frames (ORFs) Rh157.5 and Rh157.4 (the homologs of HCMV UL130 and UL128, respectively) from the genome of FL-RhCMV **(Fig. 1A)**. Both genes were also naturally lost from the parental 68-1 strain during tissue culture and the remaining three subunits are either essential for growth *in vitro* (gH and gL) (*45*) or for *in vivo* infection (Rh157.6 (UL131A)) (*46*). Since 68-1 also lacks CXC chemokine-like ORFs homologous to HCMV UL146, we deleted the six ORFs related to UL146 in a separate recombinant to evaluate the contribution of PC-deficiency versus UL146 family-deficiency during primary infection. As expected, deletion of these genes did not negatively affect *in vitro* replication in fibroblasts **(sup. Fig. 1A)**. However, PC-deleted RhCMV displayed impaired entry into epithelial cells compared to FL-RhCMV as shown by immunofluorescence (**sup. Fig. 1B**). These data are consistent with previous findings indicating that the PC enables more efficient virus entry into epithelial cells in vitro (*47, 48*).

**Figure 1.**
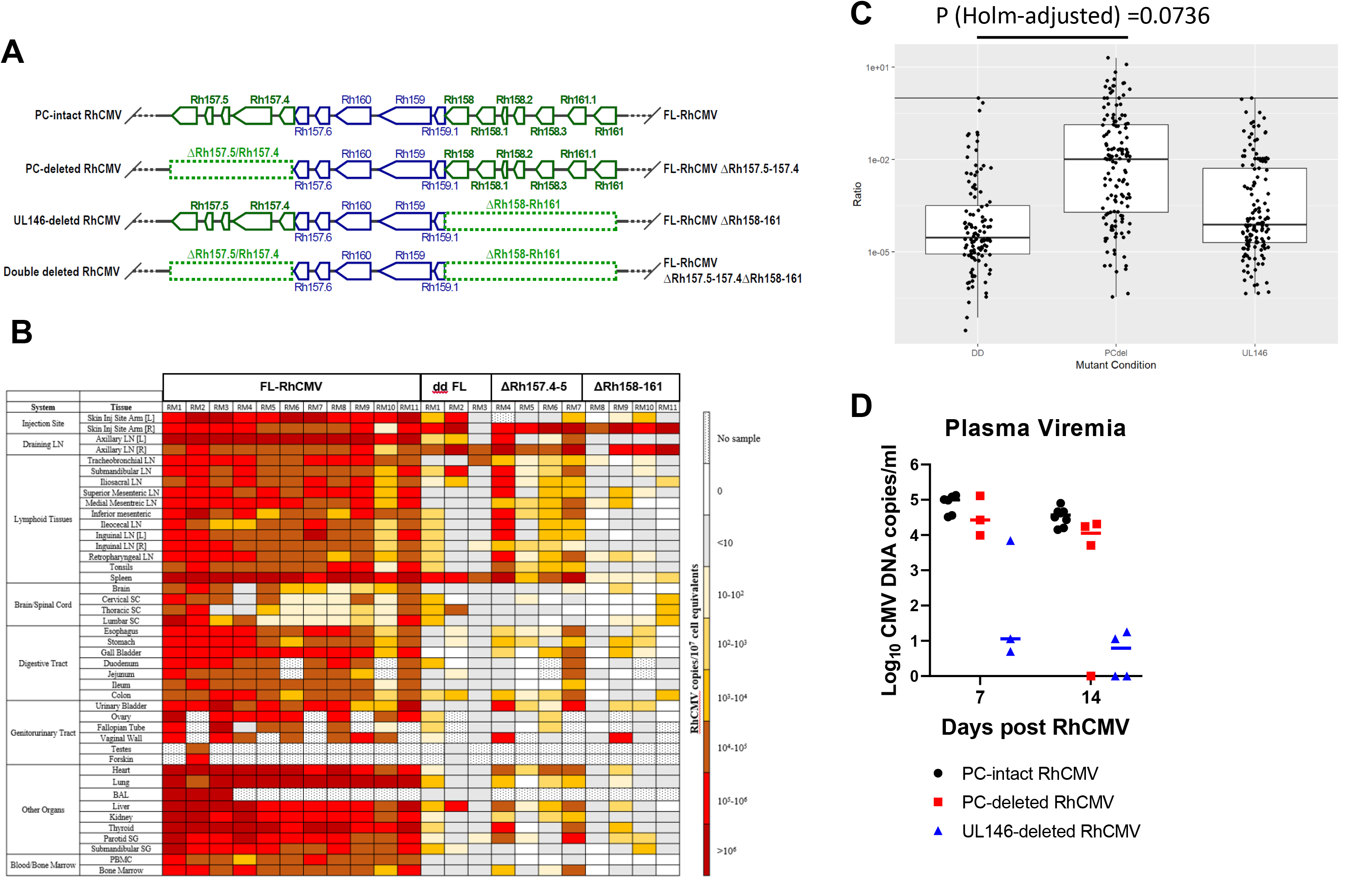
Reduced dissemination of PC-deleted and UL146-deleted RhCMV. (A) Schematic presentation of deletions in the FL-RhCMV clone. FL-RhCMV has a wildtype configuration with an intact PC. “PC-deleted” RhCMVΔRh157.5-157.4 was derived from FL-RhCMV by deleting Rh157.5 (HCMV UL128 homolog) and Rh157.4 (HCMV UL130 homolog). “UL146-deleted” RhCMV ΔRh158-161 was derived from FL-RhCMV by deleting the genes Rh158 to Rh161. “double-deleted (dd)” FL-RhCMV ΔRh157.4-157.5 ΔRh158-161 was deleted for both gene regions. (B) Tissue genome copy numbers of FL-RhCMV and deletion viruses. The Rh13.1 ORF was replaced in all recombinants with previously described heterologous inserts as immunological and PCR markers. RM1-RM3 were co-inoculated with 10^7^ PFU of FL-RhCMV (SIVgag) and dd FL-RhCMV (SIVretanef), RM4-7 were co-inoculated with FL-RhCMV (SIVgag) and PC-deleted RhCMV (TB6Ag), RM8-11 were co-inoculated with FL-RhCMV (SIVgag) and UL146-deleted RhCMV (SIVretanef). All RM were RhCMV seronegative. All RM were necropsied at day 14 post-infection and viral genome copy numbers per 10^7^ cell equivalents were determined in the indicated tissues using ultra-sensitive nested qPCR specific for each inserted antigen. Genome copy numbers are shown color coded using the color scheme shown in the right. Copy numbers are shown in **Sup Table 1**. (C) Genome copy numbers from (B) shown relative to the internal FL-RhCMV control. Statistical analysis of this dataset was performed using a linear mixed effects model excluding all tissues at the injection site and the nearest draining lymph nodes to detect significant differences in dissemination. The boxplot shows the ratio of mutant virus to the wild type virus count (after adding 1 to both counts to avoid division by zero) from the same tissue and the same animal. The solid black line indicates a ratio of 1. Analysis results are shown in **Sup Table 2**. (D) Blood viremia in PC-intact RhCMV compared to PC-deleted and UL146-deleted RhCMV. Blood samples from RM4-11 were collected 7- and 14-days post inoculation. Viral DNA copy numbers were detected by quantitative real-time and nested PCR targeting the inserted heterologous antigens. Data shown are the mean RhCMV copy numbers per 10^7^ cell equivalents of 10 individual replicates in plasma. RM7 and RM11 didn’t have blood samples collected at day 7, so only 3 RMs viral DNA copy numbers were shown at day 7.

We previously demonstrated that FL-RhCMV deleted for both the PC-subunits Rh157.5 and Rh157.4 and all six members of the UL146 family showed reduced dissemination from the initial skin injection site after s.q. inoculation of CMV-naïve RM (*33*). Furthermore, these deletions enabled RhCMV to elicit MHC-E and MHC-II-restricted CD8+ T cells instead of conventionally MHC-Ia-restricted responses (*35*), as previously reported for SIV vaccines based on RhCMV 68-1 (*34, 49*). To determine whether the reduced dissemination of the double-deleted (dd) RhCMV was due to the lack of a functional PC or due to the deletion of the UL146 family, or both, we compared viral dissemination of dd or single-deleted deletion mutants with co-inoculated FL-RhCMV in CMV naïve RM. As reported previously (*33*), high levels of FL-RhCMV genome copy numbers were detected in most tissues in all RM at 14 days post-inoculation whereas dd RhCMV was largely confined to the injection site and the nearest draining lymph nodes **(Fig. 1B-C, sup. Table 1)**. Interestingly, single deletion of the PC-components alone resulted in substantial dissemination from the skin injection site throughout the body in some RM **(Fig. 1B-C, sup. Table 1)**. In contrast, the deletion mutant lacking the UL146 family lost the ability to effectively spread past the skin injection site and the nearest draining lymph nodes very similar to dd RhCMV **(Fig. 1B-C, sup. Table 1)**. Similarly, only limited reduction of plasma viremia was observed on day 7 and 14 post infection with PC-deleted RhCMV whereas plasma viremia levels of UL146-deleted RhCMV family were severely reduced at day 7 and 14 post infection **(Fig. 1D)**. These results suggest that both the PC and the UL146 family contribute significantly **(sup. Table 2**) to the dissemination defect observed for the dd RhCMV recombinant. However, the viral CXC chemokines displayed a surprisingly strong contribution to this phenotype, even more than the PC.

### Detection of transplacental transmission in a high-risk cCMV model

CD4+ T cell responses are critical for the control of HCMV replication and, by extension, the development of morbidity in immunocompromised patients (*50–53*). We previously developed a high-risk cCMV transmission model in seronegative RM by depleting maternal CD4+ T cells (CD4-depleted) one week prior to i.v. RhCMV inoculation. This depletion increases RhCMV plasma viremia and viral shedding in maternal bodily fluids during the acute phase of the primary CMV infection, resulting in a robust and reproducible cCMV transmission and a high rate of fetal loss (*54*).

To investigate whether the PC is required for RhCMV vertical transmission in this high-risk transmission model we assigned six seronegative, pregnant dams into two groups (N=3 for each group) and depleted maternal CD4+ T cells using a rhesus IgG1 recombinant anti-CD4 monoclonal antibody (CD4R1) **(Sup Fig. 2A-F)**. FL-RhCMV was used in the control group and designated as PC-intact RhCMV in the following studies. Each animal was infected i.v. with either 10^6^ PFU of PC-intact or PC-deleted RhCMV at the end of the first trimester **(Fig. 2A)**. Two dams inoculated with the PC-deleted RhCMV experienced fetal loss at two weeks post inoculation. These two dams, as well as one dam from the PC-intact group, required euthanasia due to decompensation resulting from CMV dissemination. While an increased rate of spontaneous abortions in CD4-depleted dams after primary RhCMV infection is consistent with previous results from our group(*55*), this outcome was unexpected after infection with the PC-deleted RhCMV, considering the significantly reduced dissemination observed *in vivo* **(Fig. 1)**. To ensure that the transmitted virus was indeed lacking the PC, we collected maternal fluid and fetal tissue samples, isolated the DNA, and performed quantitative PCRs (qPCRs) targeting the Rh157.5 (UL128), Rh157.4 (UL130), and Rh157.6 (UL131A) ORFs encoding different neighboring PC subunits, of which only Rh157.6 should be retained in the genome. As expected, only primer/probe pairs targeting Rh157.6 but not Rh157.5 and Rh157.4 amplified their target DNA sequences **(Sup Fig. 3A-C)** demonstrating that the PC-deleted virus had indeed crossed the placenta resulting in a congenital infection.

**Figure 2.**
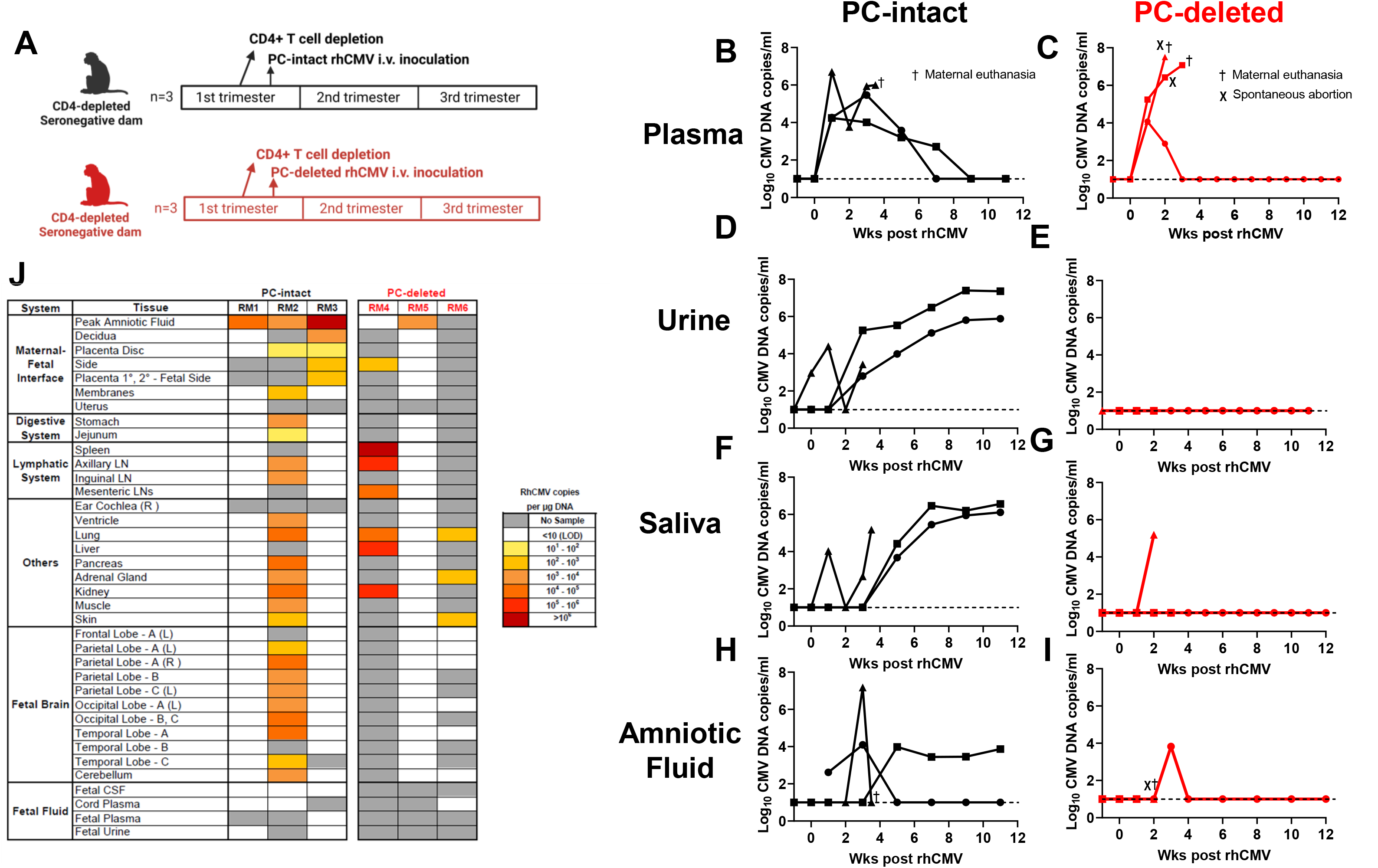
CD4-depleted dams inoculated with PC-deleted RhCMV showed similar peak viremia and transmission rate but reduced viral shedding in urine and saliva. (A) Experimental scheme for CD4-depleted RMs. Seronegative dams were CD4+ T cell-depleted 1 week prior to inoculation with 1 × 10^6^ PC-intact or 1 × 10^6^ PC-deleted RhCMV at the end of the 1st trimester (N=3 each group, respectively). Maternal samples (blood, urine, saliva, amniotic fluids) were collected every other week throughout the study period and amniotic fluid weekly from weeks 2-4 post-inoculation then every other week (6-12 weeks) until hysterotomy. Fetal tissue samples were collected at hysterotomy. Two dams were euthanized due to decompensation resulting from disseminated RhCMV at 2 weeks (spontaneous abortion) or 3 weeks (early hysterotomy) post-inoculation. (B-I) RhCMV copy numbers in maternal plasma (B,C), urine (D,E), saliva (F,G), and amniotic fluid (H,I) were estimated by real-time PCR targeting gB. Data shown as the mean RhCMV copy number of 3 individual replicates. The dotted line indicates the assay limit of detection (10 copies per ml of fluids). Black icons: dams inoculated with PC-intact RhCMV; red icons: dams inoculated with PC-deleted RhCMV. †: dam euthanasia due to poor maternal health. Χ: spontaneous abortion. (J) RhCMV copy numbers per µg DNA in maternal-fetal interface and fetal tissues were estimated by real-time PCR targeting gB. Each cell value is the mean RhCMV copy number of 3 individual replicates. Viral DNA copies are color-coded. Gray: no sample collection.

CD4-depleted dams i.v. inoculated with PC-deleted RhCMV demonstrated plasma viremia levels similar to or slightly above levels observed after infection with PC-intact RhCMV **(Fig. 2B-C**). Conversely, acute infections with the PC-intact RhCMV resulted in higher levels of viral genome copy numbers in urine, saliva, and amniotic fluid samples in all dams compared to acute infections with the PC-deleted FL-RhCMV which was barely detectable (**Fig. 2D-I)**. These results are based on a very small sample size as two out of three PC-deleted RhCMV-infected dams aborted shortly after virus inoculation, likely prior to the onset of detectable viral DNAemia in saliva, urine, and amniotic fluid. Still, the data suggests that deletion of the PC renders RhCMV impaired for dissemination, consistent with results observed after s.q. inoculation. In human clinical patients, a cCMV infection is generally determined prenatally through a positive diagnostic PCR assay of an amniotic fluid sample (*56, 57*). In our model, we were able to detect RhCMV in amniotic fluid from both infection groups, implying that the PC-deleted RhCMV can be transmitted vertically in CD4-depleted dams **(Fig. 2H, I)**. To determine whether we could detect congenital transmission not only in the amniotic fluid but also in fetal tissues, we performed quantitative PCR (qPCR) on DNA isolated from all pregnancies where fetal tissues were available **(Fig. 2J)**. RhCMV genomes were detected at the maternal-fetal interface or in fetal tissues in two out of three fetuses from dams inoculated with PC-intact or PC-deleted RhCMV, respectively. Combining results from the amniotic fluid and fetal tissues, we confirmed vertical transmission in all animals from both groups. Taken together, these results confirm that the PC-deleted RhCMV, despite demonstrating reduced dissemination during acute infection upon s.q. or i.v. inoculation, is still able to cross the placenta, suggesting that the RhCMV PC is not required for cCMV infection in CD4-depleted dams.

### PC-intact and -deleted RhCMV infection leads to similar cCMV transmission rates in immunocompetent dams

We next investigated whether the PC is required for vertical transmission in CMV seronegative, immunocompetent dams. We divided the RM into two groups (N=6 for each group) and inoculated the control group i.v. with a mixture of 1×10^6^ PFU each of the low passaged RhCMV isolate UCD52, which had previously been shown to establish congenital infection, and PC-intact RhCMV. The test group received 1×10^6^ PFU of PC-deleted RhCMV i.v., with all inoculations performed at the beginning of the 2nd trimester **(Fig. 3A)**. Dams from both groups reached peak plasma viremia within 2-3 weeks post inoculation, with the PC-deleted group displaying similar or slightly higher average peak viremia levels, yet faster control of plasma viremia than the PC-intact group. However, no statistically significant differences were observed comparing peak viremia levels between groups (P value=0.937, **Table 1**) **(Fig. 3D, E)**. Consistent with findings in the CD4-depleted dams, PC-deleted RhCMV DNA copies were rarely detectable in urine and saliva samples **(Fig. 3F, G)**. As discussed above, these results are consistent with PC-deficient RhCMV manifesting reduced propensity for dissemination.

**Figure 3.**
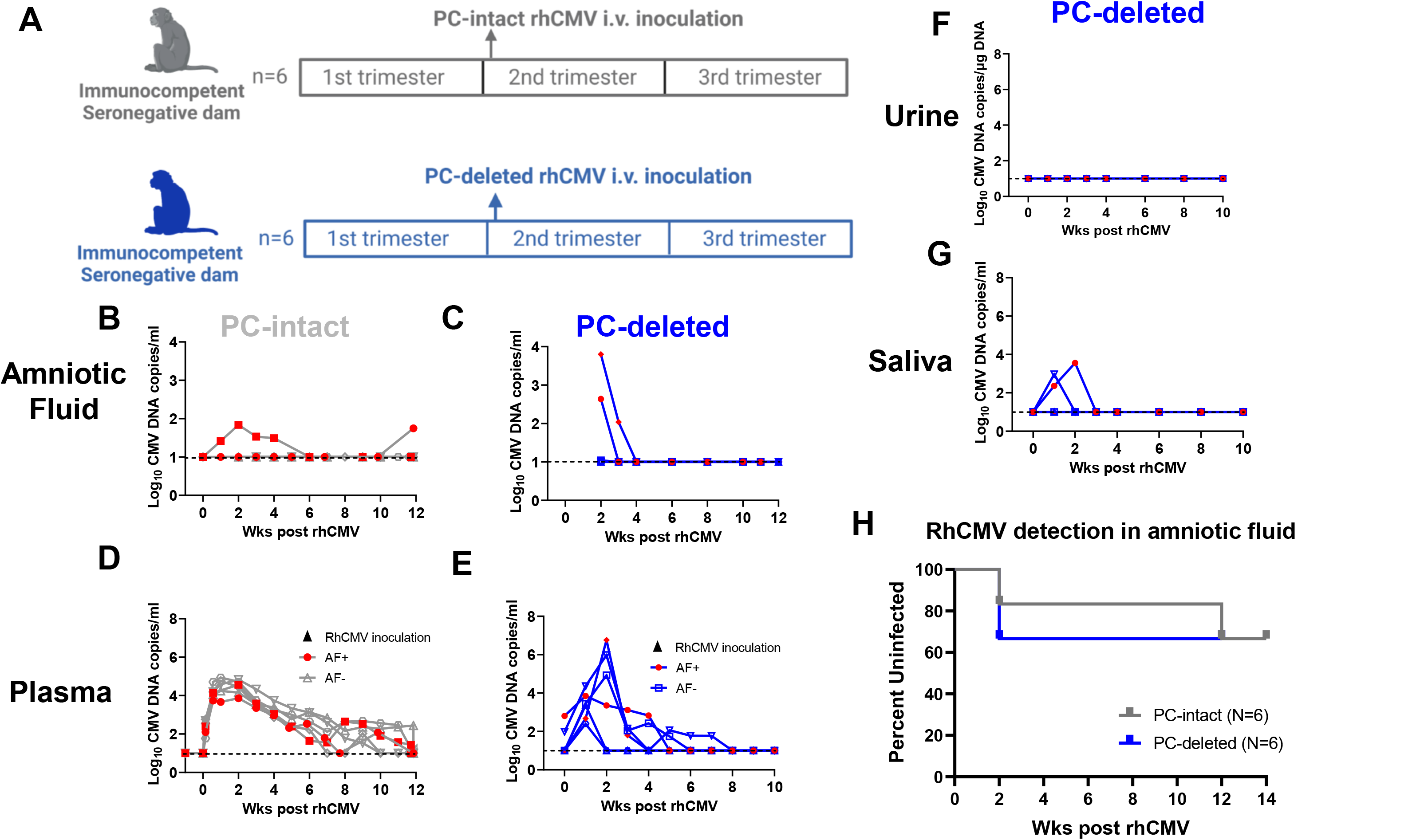
Immunocompetent dams inoculated with PC-deleted RhCMVshowed faster viremia clearance, similar shedding in AF, and reduced shedding in urine and saliva. (A) Experimental scheme of immunocompetent RMs. Seronegative dams were inoculated with 2 × 10^6^ PC-intact RhCMV (1 × 10^6^ UCD52 clinical isolate, 1 × 10^6^ FL RhCMV) or 1 × 10^6^ PC-deleted RhCMV at the end of the 1st trimester (N=6 each group, respectively). Maternal samples (blood, urine, saliva) were collected weekly throughout the study period and amniotic fluid weekly from weeks 2-4 post-inoculation then every other week (6-12 weeks) until hysterotomy. Fetal tissue samples were collected at hysterotomy. (B-G) RhCMV copy numbers in maternal amniotic fluid (B, C), maternal plasma (D, E), urine (F), or saliva (G) were estimated by real-time PCR targeting IE Exon 4. Maternal samples were collected from dams inoculated with PC-intact (B, D) or PC-deleted RhCMV (C, E, F-G). Data shown as the mean RhCMV copy number of 6 or 12 individual replicates. Fluid samples were considered positive if RhCMV DNA was detected in 2 or more out of 6 or 12 replicates. The dotted line indicates the assay limit of detection (10 copies per ml of fluid). Gray open icons with gray line: dams inoculated with PC-intact RhCMV, no virus detected in AF; red closed icons with gray line: dams inoculated with PC-intact RhCMV, virus detected in AF; Blue open icons with blue line: dams inoculated with PC-deleted RhCMV, no virus detected in AF; red closed icons with blue line: dams inoculated with PC-deleted RhCMV, virus detected in AF. A Wilcoxon Rank Sum Test was used to test for differences in peak plasma viremia by virus type. (H) Time of RhCMV detection in amniotic fluid shown by the survival curve.

**Table 1.** Statistical analysis for peak viremia and maximum % ADCP response in immunocompetent dams using AUC analysis.

| Variable | P value* |
| --- | --- |
| Peak plasma viremia (RhCMV copies/ml) | 0.937 |
| Maximum %ADCP against PC-intact RhCMV whole virion | 0.132 |
| Maximum %ADCP against PC-deleted RhCMV whole virion | 0.589 |

\*Unadjusted P value from Wilcoxon Rank Sum Test

We next evaluated the RhCMV DNA genome copy numbers in amniotic fluid over time following acute infection by qPCR. Overall, two out of six immunocompetent dams from each group had a positive amniotic fluid PCR result at least once over the course of the study **(Fig. 3B, C)**. Interestingly, for one dam inoculated with the PC-intact RhCMV mixture, we started detecting viral genomic DNA in the amniotic fluid late in the 3rd trimester at approximately 12 weeks post inoculation, while all other transmitting dams had positive amniotic fluid results at earlier time points at approximately two weeks post virus inoculation **(Fig. 3B, H)**. In addition to the amniotic fluid, we determined the RhCMV viral load in fetal tissues and placenta collected from animals in the PC-deleted group **(Sup Fig. 7)**. Unlike the high risk cCMV model, no RhCMV DNA was detected in any of the fetal tissue samples in PC-deleted or PC-intact-infected dams from the immunocompetent group. This result emphasizes that maternal immunity is able and essential to limit dam to fetus cCMV transmission. Nonetheless, low RhCMV DNA copies were observed in the placenta of four out of six animals from the PC-deleted group, suggesting that the PC-deleted RhCMV transiently replicated or traversed the placenta without transplacental transmission to the fetus **(Sup Fig. 7A)**. Consistent with this interpretation, hematoxylin and eosin (H&E) staining of placental tissue sections showed no observable pathology or signs of RhCMV infection or replication **(Sup Fig. 7B-C)**. Overall, our data demonstrate that the PC is not required for cCMV infection of CD4-depleted or fully immunocompetent, RhCMV seronegative dams.

### RhCMV-specific IgG binding and functional antibody responses elicited by PC-deleted and PC-intact RhCMV in CD4-depleted and immunocompetent dams

PC-specific antibody responses have been investigated in vertically CMV-transmitting and non-transmitting pregnant women (*58*), as well as in HCMV V160 vaccine study (*59*). Our high- and standard-risk cCMV RM model provided an opportunity to study humoral immune responses elicited in the presence and absence of the PC expressed on the surface of RhCMV virions.

We previously reported that CD4-depleted dams mounted a delayed humoral immune response against RhCMV compared to fully immunocompetent animals (*55*). Consistent with our prior results, a two to three-week delay in the development of RhCMV-IgG binding and functional humoral immune responses including NAb and antibody-dependent cellular phagocytosis (ADCP) was observed in CD4-depleted dams **(Sup Fig. 6)** (we did not perform statistics due to the small number of animals in each group). As expected, we observed reduced IgG binding to soluble, recombinant PC and lower titers of antibodies neutralizing PC-dependent epithelial cell infection in maternal plasma collected from the CD4-depleted group infected with the PC-deleted RhCMV compared to the group infected with the PC-intact virus **(Sup Fig. 6G-H, K-L).** In contrast, IgG binding to PC-intact and PC-deleted RhCMV virions and to recombinant RhCMV gB were similar in both groups as were NAb titers measured by fibroblast infection and ADCP responses **(Sup Fig. 6A-F, I-J, M-P)**.

We next compared the RhCMV-specific IgG responses elicited in immunocompetent, seronegative dams infected with PC-intact or PC-deleted RhCMV **(Fig. 4A-D)**. Dams inoculated with PC-intact RhCMV mounted an IgG binding response to PC-intact RhCMV virions two to three weeks earlier compared to dams inoculated with PC-deleted RhCMV (P value=0.0195, FDR P value=0.0438, **Table 2**). As expected, the maximal IgG response against PC-intact RhCMV virions was higher in the PC-intact group compared to almost all dams in the PC-deleted group at later time points during infection.

**Figure 4.**
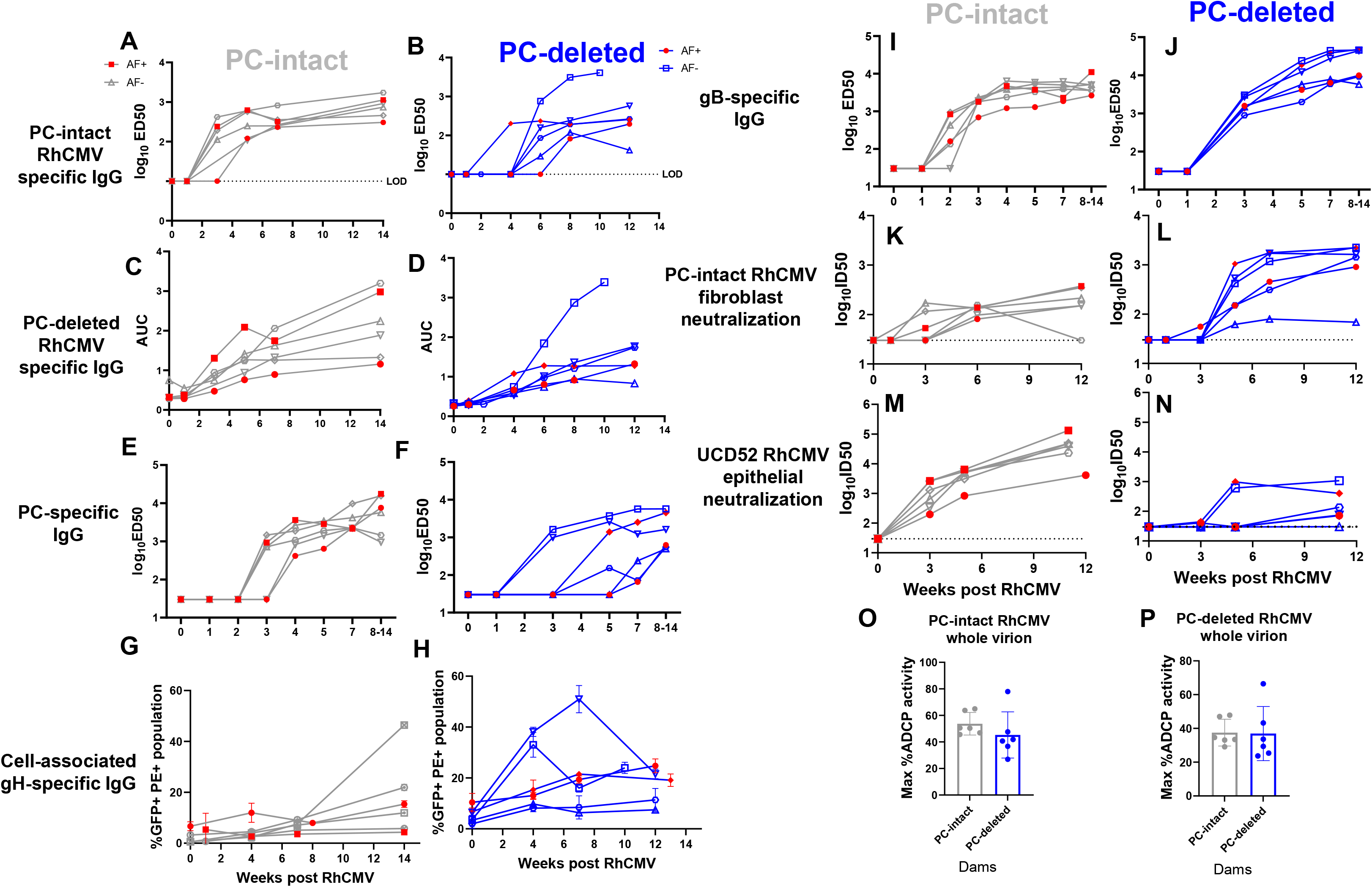
IgG binding, neutralizing antibody titers, and antibody-dependent phagocytosis response in immunocompetent dams. (A-J) Rhesus plasma IgG binding kinetics to RhCMV whole virions and glycoproteins post virus inoculation. Rhesus plasma IgG binding to PC-intact RhCMV whole virions (A,B), PC-deleted RhCMV whole virions (C,D), RhCMV PC (E-F), and RhCMV gB (I-J) by ELISA. Rhesus plasma IgG binding to cell-associated RhCMV gH (G-H) by flow cytometry. Data points are shown as the ED50 or AUC of each animal at selected timepoints. (K-N) Rhesus plasma IgG neutralization on fibroblasts against PC-intact RhCMV (K, L) and on epithelial cells against UCD52 strain (M, N). Data points are shown as the ID50 of each animal at selected timepoints. The dotted line indicates the assay limit of detection. Gray open icons with gray line: dams inoculated with PC-intact RhCMV, no virus detected in AF; red closed icons with gray line: dams inoculated with PC-intact RhCMV, virus shedding detected in AF; Blue open icons with blue line: dams inoculated with PC-deleted RhCMV, no virus shedding detected in AF; red closed icons with blue line: dams inoculated with PC-deleted RhCMV, virus shedding detected in AF. (O-P) Rhesus plasma IgG maximum % ADCP activity against PC-intact RhCMV whole virions (O) and PC-deleted RhCMV whole virions (P). Data points are shown as the maximum %ADCP activity of each animal. Gray closed icons: dams inoculated with PC-intact RhCMV; blue closed icons: dams inoculated with PC-deleted RhCMV. For longitudinal data, we fit a linear mixed effects model accounting for the data at a lower bound. Covariates in the model included Time as week, early period (before week 3), and when possible, an interaction between time and early period. For max ADCC, Wilcoxon rank sum test was used.

**Table 2.**
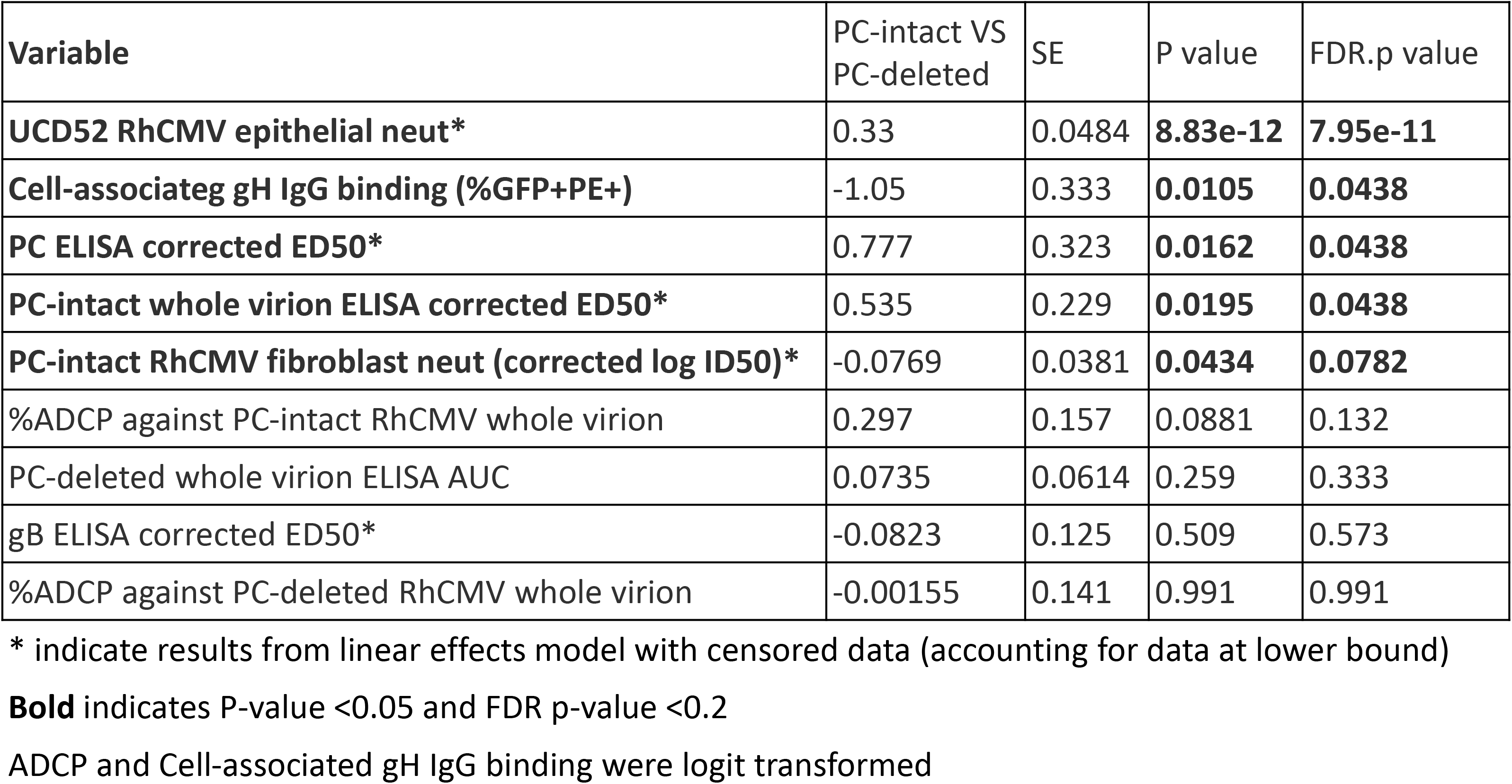
Statistical analysis for humoral responses in immunocompetent dams using the linear mixed effects model.

We next examined the RhCMV glycoprotein-specific IgG response against gB, the most widely used vaccine targets to date, and recombinant, purified PC. Dams inoculated with PC-intact RhCMV mounted a PC-specific IgG response at three to four weeks post inoculation and maintained a statistically significant higher response than the PC-deleted group throughout pregnancy (P value=0.0162, FDR P value=0.0438, **Table 2**) **(Fig. 4E-F)**. However, the kinetics of the PC-specific IgG binding response in the RM group inoculated with the PC-deleted RhCMV varied between individual animals **(Fig. 4F)**. One possible explanation for this observation could be overlapping humoral immune responses against components of the gH/gL/gO trimer with components of the (UL128/UL130/UL131A/gH/gL) pentamer as gH and gL are part of both protein complexes. Therefore, we designed a binding assay by transiently expressing gH on the surface of transfected cells to measure IgG binding of rhesus plasma derived from the control and the test group. Consistent with our hypothesis, the PC-deleted group elicited an earlier and significantly higher IgG binding response to cell-associated gH compared to the PC-intact group **(Fig. 4G-H)**. Dams from the PC-deleted group demonstrated a slightly higher IgG binding response to gB than the PC-intact group **(Fig. 4I - J)**. These findings suggest that gH and gB may become more immunodominant targets of the humoral immune responses in the absence of the PC.

*In vivo* and *ex vivo* studies have demonstrated that the PC in HCMV represents the most potent NAb target in multiple human and animal studies (*18, 60–63*). Since the PC is required for efficient infection of epithelial cell *in vitro* (*47, 48, 64, 65*), we hypothesized that dams inoculated with PC-deleted RhCMV should have a lower NAb response to infected epithelial cells compared to dams inoculated with PC-intact RhCMV. Consistent with this assumption, we indeed observed a much lower and delayed plasma NAb response to infected epithelial cells in the PC-deleted group compared to the PC-intact throughout pregnancy (P value= 8.83e-12, FDR P value= 7.95e-11, **Table 2**). **(Fig. 4M-N)**. We detected plasma Nab response to infected epithelial cells in two dams from the PC-deleted group but as described above, these responses are likely due to overlapping protein complex members between the trimer and the PC (*66*). Notably, plasma of dams inoculated with PC-deleted RhCMV demonstrated higher neutralization of fibroblast infection than those in the PC-intact group at five to six weeks post virus inoculation (P value=0.0434, FDR P value=0.0782, **Table 2**) **(Fig. 4K-L)**, which could be explained by the higher magnitude of gB- and gH-specific antibodies generated in dams inoculated with the PC-deleted RhCMV, as gB and gH/gL/gO are important for efficient entry into fibroblasts *in vitro* (*67, 68*).

We previously observed in human cohort studies that reduced cCMV transmission in seropositive mothers is correlated to a high antibody-dependent phagocytosis (ADCP) response. To investigate the role of PC-specific antibodies in mediating ADCP responses, we assessed rhesus plasma against purified virions of PC-intact and PC-deleted RhCMV **(Sup Fig. 5B-E).** Dams from both groups had a detectable ADCP response against virions from both viruses three to four weeks post inoculation, which was maintained throughout pregnancy. Comparing the maximum ADCP response, the groups showed a similar maximum ADCP response against PC-deleted RhCMV virions, while the maximum ADCP response against PC-intact RhCMV virions was slightly higher in the PC-intact virus-infected group **(Fig. 4O-P)**. However, no statistical difference was observed, suggesting PC-specific antibodies make a limited contribution to ADCP compared to non-PC targets target in our model.

## Discussions

Progress in the field of HCMV vaccine development has unfortunately been sluggish over the past decades. Although significant effort has been invested into novel vaccine concepts, no approach has demonstrated high enough efficacy to warrant licensure (*11*). The most successful clinical trials to date involved the use of an MF59-adjuvanted glycoprotein B (gB) subunit vaccine that proved to be partially protective against virus acquisition in CMV naïve women of childbearing age (*69, 70*). Alterations to the examined concept leading to improved outcomes have been proposed and largely comprise the addition of further immunogenic antigens, namely the PC (*11*). While the immunogenic potential of this protein complex has been confirmed in various preclinical animal models (*15, 71*), it is still unclear whether the strong humoral immune responses against the PC have the potential to protect the fetus from cCMV infection. Our study was meant to fill this knowledge gap by determining whether the PC is essential for transplacental CMV transmission, which could further our understanding of fetal CMV pathogenesis and guide CMV vaccine development.

Our study is the first to investigate the role of the CMV PC in placental transmission using a highly relevant non-human primate model. We constructed a PC-deleted RhCMV by removing Rh157.4 and Rh157.5, two subunits required for PC formation, from the genome. We inoculated seronegative dams intravenously with PC-intact or PC-deleted RhCMV and examined the cCMV transmission rate as well as the humoral responses elicited by RhCMV. PC-deleted RhCMV was transmitted to the fetus in both the high-risk cCMV transmission model using maternal CD4-depletion, and a standard-risk cCMV transmission model in immunocompetent dams. Moreover, the RhCMV-specific humoral responses did not differ in those immunocompetent dams with virus detected in the amniotic fluid compared to non-transmitting dams. Our findings thus challenge the assumption that the PC is a requirement for placental CMV transmission and raise the question of whether PC-elicited humoral responses will provide protection against cCMV.

PC-deleted and PC-intact RhCMV was also detected in the placenta in CD4-depleted as well as immunocompetent dams. However, we did not observe placental histomorphological changes or RhCMV localization in placentas by H&E and IHC staining in dams infected with the PC-deleted or PC-intact virus **(Sup Fig. 6)**. One reason that might explain why we could not localize RhCMV in placentas is that virus DNA copies in placenta samples were very low and difficult to detect, even using the highly sensitive qPCR method optimized in our study. Viral shedding in mucosal fluids typically peaked 2 to 3 weeks post RhCMV inoculation, but then were readily controlled, especially in the PC-deleted RhCMV infections. Since we collected placentas and fetal tissues at the end of gestation (12 to 14 weeks post RhCMV inoculation), virus replication may have been controlled at this time making it extremely difficult to detect. Another reason might be that the virus does not scatter around the placenta equally. Thus, many samples from each placental disk will be required to determine the presence of virus.

PC-deleted RhCMV was detected in multiple organ systems following acute infection in non-pregnant animals, indicating wide tissue tropism. Tissues included lymph nodes, urinary bladder, heart, liver, lung, kidney, and many other tissues although PC-deleted RhCMV had lower-level viral load in tissues than the PC-intact clone **(Fig. 1B, 1C, 2J)**. Interestingly, reduced PC-deleted RhCMV viral shedding was observed in urine, and PC-deleted RhCMV was rarely detected in saliva **(Fig. 2E, 2G, 3F-G)**. These results suggest PC deletion might render the mutant attenuated for virus dissemination and shedding. Several assumptions about the role of PC in viral tropism could be validated in further studies to better understand how PC mediates CMV entry into different cell and organ targets. The host epithelial and endothelial cell surface receptors that bind to HCMV PC have been reported. These receptors include neuropilin 2 (NRP2), platelet derived growth factor receptor α (PDGFR-α), and olfactory receptor (OR14I1) (*72, 73*). Tissues with high PC-intact RhCMV viral loads might indicate higher expression of PC-binding host cell receptors or more epithelial and/or epithelial-like cell types, whereas tissues with detectable PC-deleted RhCMV viral load indicate these regions might be composed of less epithelial and epithelial-like cells where virus components except PC are involved in CMV cell entry mechanism. However, reduced viral shedding in saliva and urine suggests PC could serve as a vaccine target for lowering CMV shedding and preventing horizontal transmission **(Fig. 2G, 3G)**. This idea was previously tested in RM in assessing PC-containing vaccines, yet these studies revealed that the vaccine-elicited immunity was not able to block virus horizontal transmission after oral inoculation with UCD52 strain (*74, 75*). Notably, both PC-intact and PC-deleted RhCMV were less detectable in the brain and spinal cord **(Fig. 1B, 2J)**, suggesting specific physical barriers within these regions might be responsible to prevent CMV entry and infection.

Dams inoculated with PC-intact RhCMV demonstrated an earlier and higher IgG response against PC-intact RhCMV virions compared to dams inoculated with PC-deleted virus, consistent with the PC being an immunodominant target of virus-specific antibody responses **(Fig. 4A-B)**. Also consistent with previous findings, we observed that plasma PC-specific IgG levels correlated with NAb titers determined on epithelial cells **(Fig. 4E-F, 4M-N)** (*14–17*). Interestingly, dams from the PC-deleted group showed slightly higher gB-specific IgG binding and generated much higher NAb titers determined on fibroblast (**Fig. 4I-L)**. These results indicate a shift of immunodominant targets when the PC is not present.

Although limited animal numbers in each group greatly impacted the statistical power, it is remarkable that cCMV transmission to the amniotic fluid occurred in 2 of 6 (33%) dams in both the PC-intact and PC-deleted dam groups, closely mirroring the rate of cCMV transmission after acute infection in human pregnancy (*7*). However, it should be noted that this model employed the i.v. route of virus inoculation which might impact cCMV transmission outcomes. Although i.v. inoculation is commonly used in experimental studies of cCMV transmission (*55, 76*), natural CMV transmission is more frequently transmitted through mucosal routes in the setting of maternal infection (*77–83*). Future studies will be required to compare the infection outcomes between different maternal inoculation methods and to develop a cCMV transmission model that mimics mucosal HCMV infection during pregnancy.

Overall, our study suggests that PC is not required for placental CMV transmission in the RM model, raising the possibility that it is also dispensable in human cCMV transmission. These results raise the question of what are the virologic determinants of cCMV transmission, which could involve other entry glycoproteins such as gB or gH/gL/gO trimer or mechanisms of viral immune evasion. Our finding that the UL146-like family was highly limited in dissemination suggests that members of this family might be required for congenital infection and could potentially be targeted. Viral components required for cCMV transmission likely represent ideal targets for vaccines that seek to prevent cCMV transmission at the level of the maternal-fetal interface even after mucosal virus acquisition. Additionally, both PC and non-PC directed NAb and non-NAb responses that are higher level than that elicited by natural immunity could be considered as a goal for CMV vaccines, as placental transmission continues to occur in seropositive women, although at a lower rate compared to primary infection (*7*). The outcome of the currently enrolling HCMV gB and PC mRNA vaccine trials will reveal the ability of antibodies against these targets to block HCMV acquisition, but longer-term studies will be required to determine if the responses can block placental transmission in women who become pregnant. Thus, pre-clinical and clinical studies should further assess the virologic and immunologic determinants of cCMV transmission in order to achieve a reduction in this devastating *in utero* infection through vaccination.

## MATERIALS AND METHODS

### Ethics statement

All animal procedures conformed to the requirements of the Animal Welfare Act and protocols were approved prior to implementation by the Institutional Animal Care and Use Committee at the University of California, Davis (UC Davis) (A3433-01), Tulane University (A4499-01), and OHSU (A3304-01).

### Animals and Sample Collection

At UC Davis, normally cycling, adult female rhesus monkeys (*Macaca mulatta*) (N=12; ∼5.0-9.8 kg, ∼5-11 years of age) confirmed seronegative for RhCMV and with a history of prior pregnancy were bred according to established methods, with pregnancy identified by ultrasound (*84*). Activities related to animal care (diet, housing) were performed as per California National Primate Research Center standard operating procedures (SOPs). All fetuses were sonographically assessed to confirm normal growth and development prior to maternal interventions (e.g., CD4-depletion, RhCMV inoculation). The dams were administered ketamine hydrochloride (10 mg/kg, intramuscular-IM) or telazol (5-8 mg/kg; IM) for these and subsequent ultrasound examinations. Sonographic measurements included fetal biometrics (e.g., biparietal diameter, femur length) in addition to gross anatomical evaluations (axial and appendicular skeleton, viscera, membranes, placenta, amniotic fluid volumes), which were assessed weekly across gestation. All measures were compared to normative growth charts for rhesus monkey fetuses, and all were found to be within normal limits (*84*). Maternal samples (blood, urine, saliva) were collected weekly throughout the study period. Amniotic fluid was collected weekly under ultrasound guidance from weeks 2-4 post-inoculation then every other week (6, 8, 10) until hysterotomy (∼12 weeks post-inoculation). Fetal fluids (blood, CSF, urine) and a large range of tissues as well as the placenta, decidua, membranes, and umbilical cord were collected post-hysterotomy.

At Tulane, Indian-origin RM (N=6) were from a RhCMV-seronegative expanded specific pathogen–free (eSPF) colony housed at the Tulane Primate National Research Center (or previously at the New England Primate Research Center). The RM were maintained in accordance with institutional and federal guidelines for the care and use of laboratory animals (*85*). All dams were screened for RhCMV-specific IgM and IgG by whole virion ELISA and confirmed RhCMV-seronegative before enrolling in the study. Dams (N=6; 4-13 years of age) at the Tulane NPRC were housed with RhCMV-seronegative males and screened for pregnancy every three weeks. Successful pregnancy was confirmed by abdominal ultrasound, described previously(*86*). Gestational age was estimated by sonography, based on gestational sac size (average of 3 dimensions) and greatest length.

At approximately week 7 of gestation, 6 dams (UC Davis) in the CD4-depleted groups were administered a 50 mg/kg dose of recombinant rhesus CD4+ T cell–depleting antibody (CD4R1 clone; NIH Nonhuman Primate Reagent Resource) by i.v. infusion. The antibody preparation was infused over an ∼20-minute time period. Successful maternal CD4+ T cell depletion was confirmed by flow cytometry (*87*) **(Sup Fig. 1)**. In CD4-depleted groups, 1 × 10^6^ PC-intact or 1 × 10^6^ PC-deleted RhCMV were administered i.v. to dams 1 week post CD4+ T cell depletion, i.e., week 8 of gestation (N=3 each group). In immunocompetent groups, dams were inoculated with 2 × 10^6^ PC-intact RhCMV (1 × 10^6^ UCD52 clinical isolate, 1 × 10^6^ FL RhCMV) or 1 × 10^6^ PC-deleted RhCMV at week 8 of gestation. Maternal blood, urine (ultrasound guided cystocentesis; ∼1-2 ml), saliva, and amniotic fluid (∼1 ml under ultrasound guidance) were collected on a weekly to bi-weekly basis until hysterotomy near term (∼140 days gestational age; gestational weeks 19-22; term 165±10 days). A large range of fetal and placental specimens were collected as noted. Recovered tissues were obtained when possible from dams with a spontaneous abortion.

All RM housed at the ONPRC were handled in accordance with good animal practice, as defined by relevant national and/or local animal welfare bodies. The RM were housed in Animal Biosafety level (ABSL)-2 rooms. The rooms had autonomously controlled temperature, humidity, and lighting. Study RM were both pair and single cage housed. Regardless of their pairing, all animals had visual, auditory, and olfactory contact with other animals within the room in which they were housed. Single cage housed RM received an enhanced enrichment plan and were overseen by nonhuman primate behavior specialists.

Animals were only paired with one another if they were from the same vaccination group. RM were fed commercially prepared primate chow twice daily and received supplemental fresh fruit or vegetables daily. Fresh, potable water was provided via automatic water systems. To prepare RM for blood collection, monkeys were administered ketamine (∼7 mg/kg). Monkeys were bled by venipuncture (from the femoral or saphenous veins) and blood was collected using Vacutainers. Monkeys were humanely euthanized by the veterinary staff at ONPRC in accordance with end point policies. Euthanasia was conducted under anesthesia with ketamine, followed by overdose with sodium pentobarbital. This method is consistent with the recommendation of the American Veterinary Medical Association.

### Cell culture

Telomerized rhesus fibroblasts (TeloRF) were described previously (*88*). Primary embryonal rhesus fibroblasts (1RF) were generated at Oregon National Primate Research Center (ONPRC) and have been described before (*33*). Both cell lines were maintained in Dulbecco’s modified Eagle medium (DMEM, Gibco) containing 10% heat-inactivated fetal bovine serum (FBS, Corning), 25 mM N-2-hydroxyethylpiperazine-N’-2-ethanesulfonic acid (HEPES, Gibco), 2 mM L-glutamine (Gibco), 50 U/mL penicillin and 50 μg/mL streptomycin (Gibco), 50 μg/mL gentamicin (Gibco), and 100 μg/mL geneticin (Gibco). Monkey kidney epithelial (MKE) cells were cultured in DMEM-F12 (Gibco) supplemented with 10% FBS, 25 mM HEPES, 2 mM L-glutamine, 1 mM sodium pyruvate (Gibco), 50 U/ml penicillin and 50 μg/ml streptomycin, and 50 μg/ml gentamicin. Rhesus retinal pigment epithelial (RPE) cells were a kind gift from Dr. Thomas Shenk (Princeton University, USA) and were maintained in a 1:1 mixture of DMEM and Ham’s F12 nutrient mixture (Thermo Fisher) with 5% FBS, 1mM sodium pyruvate, and nonessential amino acids. 293T cells used for transfection were maintained in DMEM supplemented with 10% FBS, 25 mM HEPES, 50 U/mL penicillin and 50 μg/mL streptomycin. THP-1 cells (ATCC) were maintained in RPMI 1640 medium (Gibco) supplemented with 10% FBS.

### RhCMV gH plasmid design

To measure IgG binding to RhCMV gH expressed on cell surface, we designed a plasmid using the concept described previously(*89*). In plasmid design, gH ectodomain (NCBI accession number: MN437483) was conjugated with an extracellular HA tag right after the signal peptide and with a human CD4 transmembrane tail to replace the gH transmembrane and cytosolic domains. HA tag was included to indicate gH expressed on cell surface, and human CD4 transmembrane (TM) tail was reported to increase protein transportation intracellularly to cell surface(*89*). The signal peptide and transmembrane domain of gH were predicted by SignalP6.0 and TMHMM2.0, respectively (*90, 91*). A linker sequence (GGGGS) between HA tag and gH-CD4TM insert was included to prevent structural interference(*92*). The insert was cloned into the pcDNA3.1(+)-P2AeGFP plasmid with NheI and BamH1 restriction digest sites.

### RhCMV strains and isolates

The 68–1 RhCMV BAC (*93*) has been characterized extensively and we recently reported the construction of FL-RhCMV, the only clonal RhCMV BAC containing a full genome (*33*). Both viruses were derived via transfection of BAC DNA into primary rhesus fibroblasts using Lipofectamine 3000 (Thermo Fisher). Full cytopathic effect (CPE) was observed after 7–10 days and the supernatants were used to generate viral stocks. UCD52 and UCD59 RhCMV have been continuously passaged on MKE cells to maintain their PC. Seed stocks generated on MKE cells were used to infect primary rhesus fibroblasts at an MOI: 0.01. Cell culture on fibroblasts was limited to a single round to avoid the acquisition of tissue culture adaptations. After infections progressed to ∼90% CPE, supernatant and cells were collected and centrifuged at 6000 × g for 15 minutes at 4 ℃. The supernatant was passed through a 0.45 μm filter and was subsequently centrifuged at 26,000 × g for 2 hours at 4 ℃. The supernatant was decanted, and the virus pellet was resuspended and washed in ∼ 20 ml cold 1X PBS. The virus was pelleted by ultracentrifugation at 72,000 × g (Rotor SW41Ti at 21,000 rpm) for 2 hours at 4 ℃, which was then repeated once more. Finally, the supernatant was decanted, and the remaining viral pellet was thoroughly resuspended in ∼1–2 ml of cold 1X PBS. The viral stock was stored in 125 μl aliquots at -80°C until final use.

### BAC recombineering using *en passant* homologous recombination

Recombinant RhCMV clones were generated by *en passant* mutagenesis, as previously described for HCMV (*94*), and adapted by us for RhCMV (*33*). This technique allows the generation of “scarless” viral recombinants, i.e., without containing residual heterologous DNA sequences in the final constructs. The homologous recombination technique is based on amplifying an I-SceI homing endonuclease recognition site followed by an aminoglycoside 3-phosphotransferase gene conferring kanamycin resistance (KanR) with primers simultaneously introducing a homology region upstream and downstream of the selection marker into the intermediate BAC cloning product. As *en passant* recombinations are performed in the GS1783 E-coli strain that can be used to conditionally express the I-SceI homing endonuclease upon arabinose induction (*94*), expression of the endonuclease with simultaneous heat shock induction of the lambda (λ) phage derived Red recombination genes will lead to the induction of selective DNA double strand breaks with subsequent scarless deletion of the selection marker. The immunologically traceable markers used in the study, namely the SIV-mac239 GAG, RevTetNev (retanef) and the Mtb Erdman strain derived TB6Ag fusion protein, have been described before (*95, 96*). To introduce these antigens into the Rh13.1 ORF of the FL-RhCMV backbone, we first introduced a homology region flanking an I-SceI site and a KanR selection marker into the selected inserts. We then amplified the transgenes by PCR and recombined the entire insert into the desired location in the FL-RhCMV BAC. The KanR cassette was subsequently removed scarlessly as described above. All recombinants were initially characterized by XmaI restriction digests and Sanger sequencing across the modified genomic locus. Lastly all vectors were fully analyzed by next generation sequencing to exclude off-target mutations and to confirm full accordance of the generated with the predicted full genome sequence.

### Virus growth kinetics on fibroblasts *in vitro*

Primary rhesus fibroblasts (PRF, Oregon National Primate Research Center) were seeded at 50,000 cells per well in 24-well plates (Corning) and infected 24 hours later in triplicate with RhCMV recombinants at multiplicity of infection (MOI) of 0.01. Supernatants from the cell culture were collected every 3 days post infection until day 24 and stored at -80 ℃. Supernatants were tittered on primary rhesus fibroblasts seeded at 10,000 cells per well in 96-well plates (Corning) in quadruplicates. At 72 hours post infection, 96-well plates were fixed and permeabilized with 2% paraformaldehyde and 0.1% Triton X-100 in PBS. 96-well plates were blocked with 2% BSA in PBS for 30 minutes and then stained with a 1:500 dilution of mouse anti-RhCMV pp65b antibody clone (VGTI Monoclonal Antibody Core) in PBS for 1 hour followed by a 1:1000 dilution of Alexa488-conjugated anti-mouse antibodies (Invitrogen) and 3 µM DAPI (Invitrogen) in PBS for another 1 hour. 96-well plates were washed three times with PBS between all the fixing, blocking and staining steps. Infected cells were imaged on an EVOS® FL Auto Imaging System (Thermo Fisher) and cell count was performed using FIJI image processing software (*97*). The DAPI+ and pp65+ populations were estimated to determine total PRF cell number and RhCMV virion numbers, respectively.

### Nested real-time PCR

To examine the differences in dissemination between the different FL-RhCMV deletion mutants, we infected RhCMV-naïve RM s.q. with 10^7^ PFU of a deletion mutant and a FL-RhCMV control vectors in different arms. We harvested plasma from each animal at days 0, 7 and 14 and euthanized 14 days post infection and harvested tissues from which the DNA was isolated by the ONPRC Molecular Virology Support Core (MVSC) using the FastPrep (MP Biomedicals) in 1 ml TriReagent (Molecular Research Center Inc.) for tissue samples under 100 mg. Additionally, 100 μl bromochloropropane (MRC Inc.) was added to each homogenized tissue sample to enhance phase separation. 0.5 ml DNA back-extraction buffer (4 M guanidine thiocyanate, 50 mM sodium citrate, and 1 M Tris) was added to the organic phase and interphase materials, which was then mixed by vortexing. The samples were centrifuged at 14,000 × g for 15 minutes, and the aqueous phase was transferred to a clean microfuge tube containing 240 μg glycogen and 0.4 ml isopropanol and centrifuged for 15 minutes at 14,000 × g. The DNA precipitate was washed twice with 70% ethanol and resuspended in 100 to 500 μl double deionized water. For each DNA sample, 10 individual replicates (5 μg each) were amplified by first-round PCR synthesis (12 cycles of 95 ℃ for 30 seconds and 60 ℃ for 1 minute) using Platinum Taq in 50 μl reactions. Then, 5 μl of each replicate was analyzed by nested quantitative PCR (45 cycles of 95 °C for 15 seconds and 60 ℃ for 1 minute) using Fast Advanced Master Mix (ABI Life Technologies) in an ABI StepOnePlus Real-Time PCR machine. The results for all 10 replicates were analyzed by Poisson distribution and expressed as copies per cell equivalents (*98*).

### Virus entry assay into fibroblasts and epithelial cells

PRF and RPE cells were seeded in 96-well plates at 12,000 cells per well. The next day, cells were infected with serial dilutions of either FL-, double-deleted, PC-deleted, or UL146-deleted RhCMV at a 1:3 dilution scheme. At 72 hours post infection, cells were fixed with methanol for 20 minutes at -20°C. After three washes with PBS, cells were stained with mouse α-RhCMV pp65b antibody clone (VGTI Monoclonal Antibody Core) for 1 hour at 37°C. Subsequently, cells were stained with Alexa488-conjugated anti-mouse antibodies (Thermo Fisher) for 1 hour at 37°C and then with DAPI (Thermo Fisher) for 10 minutes at room temperature. Finally, images were acquired using EVOS® FL Auto Imaging System (Thermo Fisher) and analyzed using ImageJ software. Infection rates were depicted as ratios of focus forming units per mL (FFU/mL) in RPE cells versus in fibroblasts for each viral vector. All experiments were performed in two biological and three technical repeats.

### Viral load detection by QPCR targeting RhCMV gB or IE-1 Exon 4

DNA was extracted from maternal fluids (plasma, saliva, urine, amniotic fluid) and fetal tissues as previously described (*32*) or manually using the QIAmp DNA minikit (Qiagen). RhCMV DNA copies were determined by QPCR targeting gB (gB QPCR) (*99*) for CD4-depleted animals and targeting IE-1 Exon 4 for immunocompetent animals(*100*). We changed the QPCR target from gB to IE-1 exon 4 after we ran a side-by-side comparison and found out QPCR targeting IE-1 exon 4 is more sensitive to determine RhCMV DNA copies. A standard gB and IE plasmids were included in each assay to interpolate the number of viral DNA copies/ml for each sample. All the DNA extracted from dams and fetuses in the CD4-depleted group were run in triplicates. For immunocompetent dams and fetuses, amniotic fluid DNA was run in 12 replicates, placenta DNA was run in 18 replicates, and all the other samples were run in 6 replicates.

### qPCR assays determining viral genome copy numbers using primer/probes specific for Rh157.5, Rh157.4 and Rh157.6

qPCR assay was performed using primers and probes specific to the pentameric glycoprotein complex (PC) subunit genes Rh157.5 (UL128), Rh157.4 (UL130), and Rh157.6 (UL131A) as described previously (*35*). Briefly, 5 µL of each isolate sample were added to 10 µL of TaqMan Fast Advanced Master Mix solution (Thermo Fisher) with the corresponding primers and probes. QPCR reactions were performed using QuantStudio 7 Flex Real-Time PCR Systems (Applied Biosystems) and data was collected using the QuantStudio Real-Time PCR Software v1.3. PC subunit gene copies were calculated as copy numbers per either 10 ng DNA isolate or 1 mL isolate and depicted as the mean of triplicate repeats (+/-SEM).

### Hematoxylin and Eosin Staining

Sections of placenta were fixed in 10% phosphate-buffered formalin, embedded in paraffin, then sectioned at 5-6 µm and stained with hematoxylin and eosin (H&E) according to established protocols (*101*). Briefly, sections were deparaffinized in xylenes (3 × 5 min each), rehydrated in a series of graded ethanol (100, 90, 70, and 50%; 2 min each), then stained for 90 seconds in Harris Modified Hematoxylin (Fisher Chemical). After a brief tap water rinse, sections were stained for 90 seconds in Eosin Y (Fisher Chemical) then dehydrated in a series of graded ethanol rinses. The sequence and timing of graded ethanols was as follows: 50% (6 quick dips), 70% (2 × 30 sec), 90% (2 × 1 min) 100% (2 × 3 min) and xylenes (2 × 5 min each). A coverslip was applied with Cytoseal mounting medium (Richard Allan Scientific).

### Rhesus plasma IgG binding to RhCMV whole virions and RhCMV proteins by ELISA

To estimate the rhesus plasma IgG binding to RhCMV whole virions, 384-well clear-bottom ELISA plates (Corning) were coated overnight at 4℃ with 120 pfu PC-intact or PC-deleted RhCMV in 0.1M Carbonate Buffer. The coating plates were washed once with the washing buffer (PBS + 0.1% Tween 20) after overnight incubation and blocked with the assay diluent (1X PBS containing 4% whey, 15% normal goat serum, and 0.5% Tween 20) at room temperature for 1 hour. Rhesus plasma was five-fold serial diluted from 1:10 and then added to plates at room temperature for 2 hour incubation. RhCMV-specific hyperimmune globulin and seropositive plasma was included as the positive control and plate-to-plate control, while RhCMV-seronegative plasma was included as the negative control. After 2-hour incubation, the wells were washed with the washing buffer twice and incubated with horseradish peroxidase (HRP)-conjugated mouse anti-monkey IgG (Southern Biotech) at room temperature for 1 hour. After incubation, the plates were washed with washing buffer four times and later developed with the SureBlue Reserve tetramethylbenzidine (TMB) peroxidase substrate (KPL) and read at 405nm. The 50% effective dose end dilution (ED50) was calculated as the plasma dilution that resulted in a 50% reduction of the IgG binding, determined by the method of Reed and Muench (*102*). The area under the curve (AUC) was calculated by plotting normalized OD values against plasma dilutions with GraphPad Prism 9.5.0.

Rhesus plasma IgG binding to RhCMV proteins was determined following the similar approach above. 384-well clear-bottom ELISA plates (Corning) were coated overnight at 4℃ with 2 µg/ml Gb or PC (A gift from Redbiotec) in 0.1M Carbonate Buffer. The plates were washed with washing buffer once and later blocked with the assay diluent at room temperature for 1 hour. Rhesus plasma was three-fold serial diluted from 1:30 (samples collected at early timepoints) or 1:270 (samples collected at later timepoints) and then added to plates at room temperature for 1-hour incubation. Next, the plates were washed with washing buffer twice before adding the HRP-conjugated mouse anti-monkey IgG (Southern Biotech) and incubating at room temperature for 1 hour. The plates were then washed with washing buffer four times before the development with the SureBlue Reserve TMB peroxidase substrate (KPL) and read at 405nm. The ED50 value was calculated as described above (*103*).

### Glycoprotein H-transfected cell binding assay

2 × 10^6^ HEK 293T cells (ATCC) were plated in T75 flasks (Thermo Fisher) and incubated at 37℃, 5% CO_2_ overnight. 50% confluent HEK 293T cells were transfected with 6000 ng GFP-tagged RhCMV gH-CD4 TM plasmid using Effectene Transfection Reagent Kit (Qiagen) next day. A mock transfection flask was performed to validate successful GFP-tagged RhCMV gH-CD4 TM plasmid transfection. Transfected cells were incubated at 37℃, 5% CO2 for 48 hours, detached with TrypLE (Gibco), and resuspended in DMEM complete media. The viability and cell count of the resuspended cells were measured using Countess Automated Cell Counter (Invitrogen). The buffer used for antibody staining and washing is 1X PBS+ 1% FBS. 100,000 cells were plated in 96-well round bottom plate and blocked with 1:1000 Human TruStain FcX™ (Biolegend) diluted in washing buffer for 10 minutes. After blocking, the cells were washed with washing buffer once and centrifuged at 1200×g for 5 minutes to aspirate the washing buffer. Rhesus plasma samples were diluted 1:30 in washing buffer in triplicates. 200 ng/ml rhesus hyperimmune globulin was included as positive control and plate-to-plate control, while 1:30 pooled rhesus seronegative plasma was included as negative control. The diluted rhesus plasma samples and controls were incubated with the transfected cells at 37℃, 5% CO2 for 2 hours. Dead HEK293T cells were prepared by heating at 95℃ for 5 minutes as a dead cell control for the following live/dead staining procedure. After 2-hour incubation, wells were washed with washing buffer twice and later added 1:1000 Near IR live/dead staining (Invitrogen) at room temperature for 20 minutes. After incubation, wells were washed again with washing buffer twice and stained with 1:200 Mouse Anti-Monkey IgG-PE, SB108a clone (Southern Biotech) at 4℃ for 30 minutes. Simultaneously, transfected cells were stained with HA-Tag Rabbit mAb, C29F4 clone, PE Conjugate (Cell Signaling Technology) to determine the HA tag expression, indicating gH surface expression, at 4℃ for 30 minutes. The stained cells were than washed again with washing buffer twice and fixed in Formalin solution, neutral buffered, 10% (Sigma Aldrich) for 15 minutes. The fixed cells were than washed twice and resuspended in washing buffer for acquisition. Events were acquired on LSRFortessa Cell Analyzer (BD) using the HTS (high throughput screening) cassette. Data was analyzed with Flowjo software (Tree Star, Inc.). The IgG binding to cell associated RhCMV gH was measured by the % of GFP+ PE+ population among live cells **(Sup Fig. 4).** Non-specific binding of PE-conjugated Anti-Monkey IgG Fc was corrected in the analysis using bead single color control, prepared by the AbC™ Total Antibody Compensation Bead Kit (Thermo Fisher).

### Neutralization

Rhesus plasma neutralizing antibody response against fibroblasts and epithelial cells was determined against TeloRF and MKE cells, respectively. Rhesus plasma samples were heat-inactivated at 65℃ for 1 hour to inactivate proteins (e.g., complement) that might impact the neutralizing activity. 1,500 fibroblasts or 3,000 MKE cells per well were plated in 384-well black and clear-bottom plates (Corning) overnight at 37℃. On the following day, heat-inactivated rhesus plasma was three-fold serial diluted from 1:30 and mixed with PC-intact RhCMV or UCD52 strain for fibroblast or epithelial neutralization assay, respectively. The plasma-virus mixture was incubated for 1 hour and later added to the fibroblasts or epithelial cells for another 24- or 48-hour incubation, respectively, at 37℃, 5% CO_2_. After 24 or 48 hour incubation, the infected cells were fixed with 1X PBS + 10% formalin at room temperature for 15 minutes before staining. The staining buffer is 1X Dulbecco’s PBS (DPBS) + 1% FBS + 0.3% Triton X-100. Fixed cells were washed with the washing buffer once and first stained with 5 µg/ml mouse anti-RhCMV Rh151/152 monoclonal antibody in the staining buffer for 1 hour. Stained cells were than washed with PBS twice and incubated with 1:500 AF488 fluorochrome-conjugated goat anti-mouse IgG H&L (Abcam) in the staining buffer for 1 hour. After PBS wash twice, cells were stained with 1:1000 DAPI (4’,6-Diamidino-2-Phenylindole, Dihydrochloride, Invitrogen) for 10 minutes. Cells were resuspended in PBS and the total (DAPI+) and AF488+ cells were counted on the ImageXpress Pico Automated Cell Imaging System (Molecular Devices). The percent infection rate was determined by AF488+/DAPI+ cells. Neutralization titers (ID50) were calculated based on 50% reduction of the percent infected cells via the method of Reed and Muench (*102*).

### Antibody-dependent phagocytosis (ADCP) response to virions

PC-intact or PC-deleted RhCMV were conjugated with DMSO-dissolved AF647–N-hydroxy succinimide ester (Invitrogen) with constant agitation at room temperature for 1 hour. The conjugation reaction was quenched with 1 M Tris-HCl, pH 8.0. In a 96-well round-shaped plate (Corning), rhesus plasma was diluted 1:30 in 1X PBS+ 1% FBS. 100 ng/ml rhesus hyperimmune globulin was included as positive control and plate-to-plate control, while 1:30 pooled rhesus seronegative plasma was included as negative control. Rhesus plasma samples and controls were incubated with viruses at 37℃ for 2 hours. After incubation, 250,000 cells/ml THP-1 cells (ATCC) were resuspended in RPMI+10% FBS media and 200μl THP-1 cells per well were added to the sera-virus mixture. The 96-well plate was later centrifuged at 1200×g, 4℃ for 1 hour to allow spinoculation and then incubated at 37°C, 5% CO_2_ for an additional 1 hour. The cells were spun down at 1200×g for 5 minutes and washed with 1X PBS+ 1% FBS once before staining with Aqua Live/Dead stain (Invitrogen) at room temperature for 25 minutes. The stained cells were washed with 1X PBS+1% FBS twice and fixed with 1X PBS + 10% formalin at room temperature for 15 minutes. After fixation, the fixed cells were washed with 1X PBS+1% FBS twice and resuspended in 100 μl PBS for the acquisition. Events were acquired on an LSRFortessa (BD Biosciences) using the HTS (high throughput screening) cassette. Data was analyzed with Flowjo software (Tree Star, Inc.). The ADCP activity was determined by the AF647+ cells from the live cell population using the pooled rhesus seronegative plasmas negative control as the threshold **(Sup Fig. 5A)**.

### Statistical analysis

We used a linear mixed effects model to fit the log10 ratio of genome copy numbers measured for each deletion virus (PC-deleted, UL146-deleted, and dd-deleted RhCMV) to FL-RhCMV in the same tissue from the same animal, excluding the injection site and draining lymph nodes. The model included random effect for each animal, and a fixed effect for the organ system. For testing if dissemination of a deletion virus differed from FL-RhCMV, we tested if the log10 ratios are different than 0. For comparison of two deletion virus strains, we tested if the difference between the deletion RhCMV number 1 normalized to their FL-RhCMV and deletion RhCMV number 2 normalized to their FL-RhCMV was different than 0. Results are considered statistically significant if the Holm-adjusted p-value was less than 0.1 and the unadjusted p-value less than 0.05. Analysis results are shown in **(Sup Table 2)**.

In the immunocompetent dams, peak viral load and max ADCP were analyzed via a Wilcoxon rank sum test, testing for differences between the two groups. We used the coin package in R(*104*). In addition, tested the AUC out to week 11 for differences in the humoral responses **(Fig. 4)** and the two ADCP measures via Wilcoxon rank sum test (data not shown). Multiple testing correction was done via FDR, with significance assessed as an unadjusted p-value less than 0.05 and an FDR corrected less than 0.2. In addition, for each humoral response (and ADCP-data not shown), we tested the association between the response and inoculated virus over time via a repeated measures analysis with a random intercept for macaque. Covariates in the model included, time (weeks), early infection period (defined as before week 3) and the interaction (when applicable) between these two variables. Data was log10 transformed (ADCP logit transformed). If there was no data at the lower bound, we used the lme4 R package (*105*) while for data at the lower bound we utilized the lmec R package (*106–108*) to account for censored observations. Multiple correction was done as above. All analyses were performed within R (*109*).

## Acknowledgments

We thank the animal care and veterinarians at the Oregon National Primate Research Center, Tulane National Primate Research Center, and California National Primate Research Center. We thank the Molecular Virology Core at Oregon Health & Science University (Drs. Greg Dissen and Don Siess, Jeffrey Torgerson, Ashley White) for RhCMV viral load quantification. We thank support from the Biostatistics, Epidemiology and Research Design (BERD) Methods Core for the statistical analysis.

---

**Sup Figure 1.**
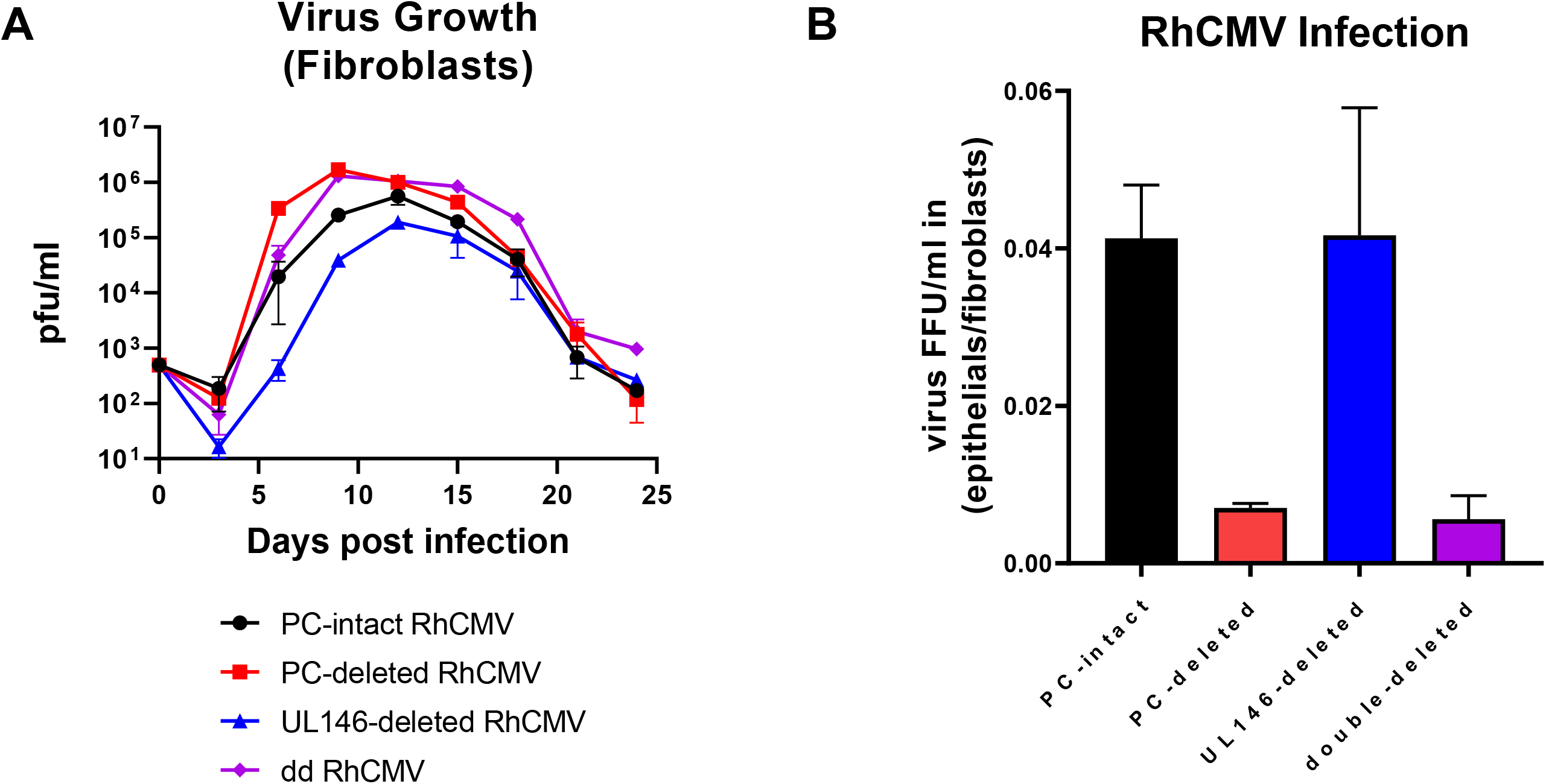
In vitro characterization of FL-RhCMV and FL-RhCMV-derived recombinants deleted for PC-subunits or the UL146 family. (A) PC-intact, PC-deleted, UL146-deleted and double-deleted RhCMV growth kinetics on primary rhesus fibroblasts *in vitro*. Data shown are the mean RhCMV numbers in fibroblast (pfu/ml) for 12 individual replicates. (B) The relative entry of PC-intact, PC-deleted and UL146-deleted RhCMV into epithelial cells compared to fibroblasts was measured using immunofluorescence assay (IFA). The epithelial/fibroblast cell entry ratio was calculated by normalizing RhCMV entering epithelial cells to RhCMV entering fibroblasts. Data shown are the mean epithelial/fibroblast cell entry ratio for 6 individual replicates.

**Sup Table 1.**
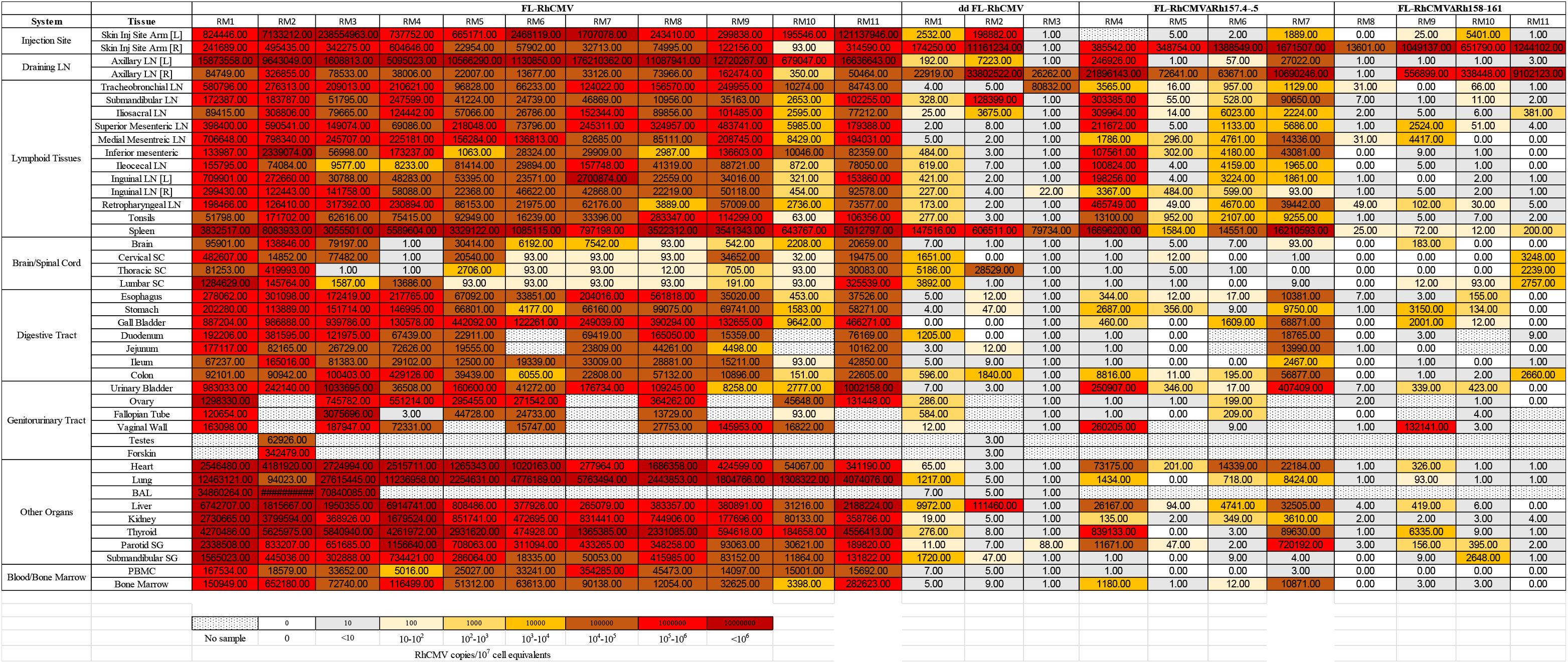
Virus dissemination of PC-deleted or UL146/147-deleted RhCMV.

**Sup Table 2.** Statistical analysis for tissue genome copy numbers of FL-RhCMV and deletion viruses using the linear mixed effects model.

| Comparison | Estimate | test_statistic | P value | Holm-Adjusted P value |
| --- | --- | --- | --- | --- |
| dd RhCMV vs FL-RhCMV | -4.86 | -8.36 | 1.53e-06 | **7.64e-06** |
| PC-deleted RhCMV vs FL-RhCMV | -2.97 | -5.7 | 4.57e-05 | **0.000183** |
| UL146-deleted RhCMV vs FL-RhCMV | -4.34 | -8.33 | 6.05e-07 | **3.63e-06** |
| dd RhCMV vs PC-deleted RhCMV | -1.89 | -2.77 | 0.0245 | **0.0736** |
| dd RhCMV vs UL146-deleted RhCMV | -0.518 | -0.759 | 0.47 | 0.47 |
| PC-deleted RhCMV vs<br>UL146-deleted RhCMV | 1.37 | 2.17 | 0.0622 | 0.124 |

**Bold** indicates P-value <0.05 and Holm-adjusted P-value <0.1

**Sup Figure 2.**
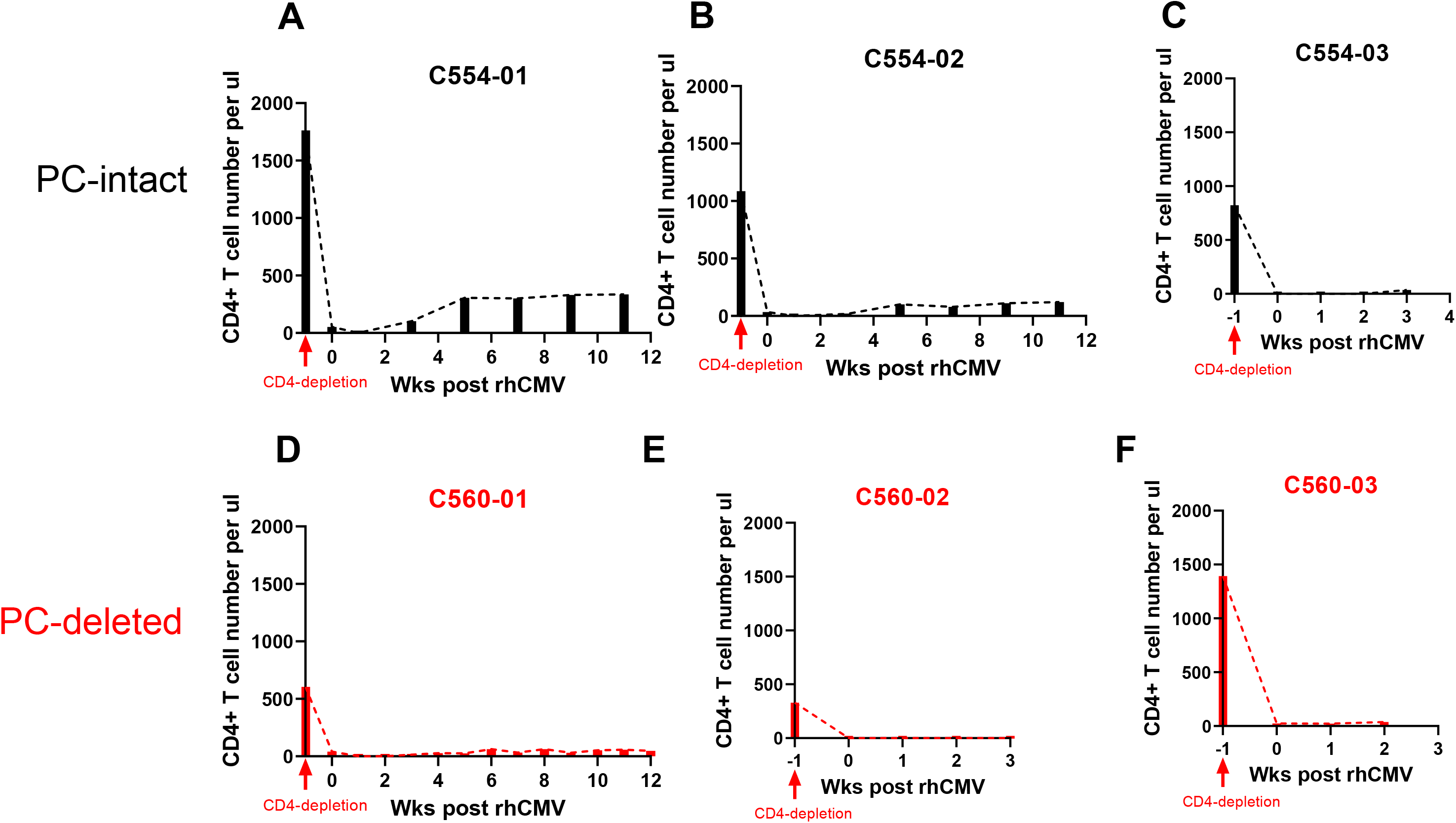
CD4+ T cells were successfully depleted post CD4R1 treatment. CD4+ T cell count of RMs was determined by flow cytometry. Maternal CD4+ T cell depletion was observed in dams inoculated with PC-intact RhCMV (A-C) or PC-deleted RhCMV (D-F). Data shown as the CD4+ T cell count in per µl blood. The dotted line indicates the CD4+ T cells kinetics.

**Sup Figure 3.**
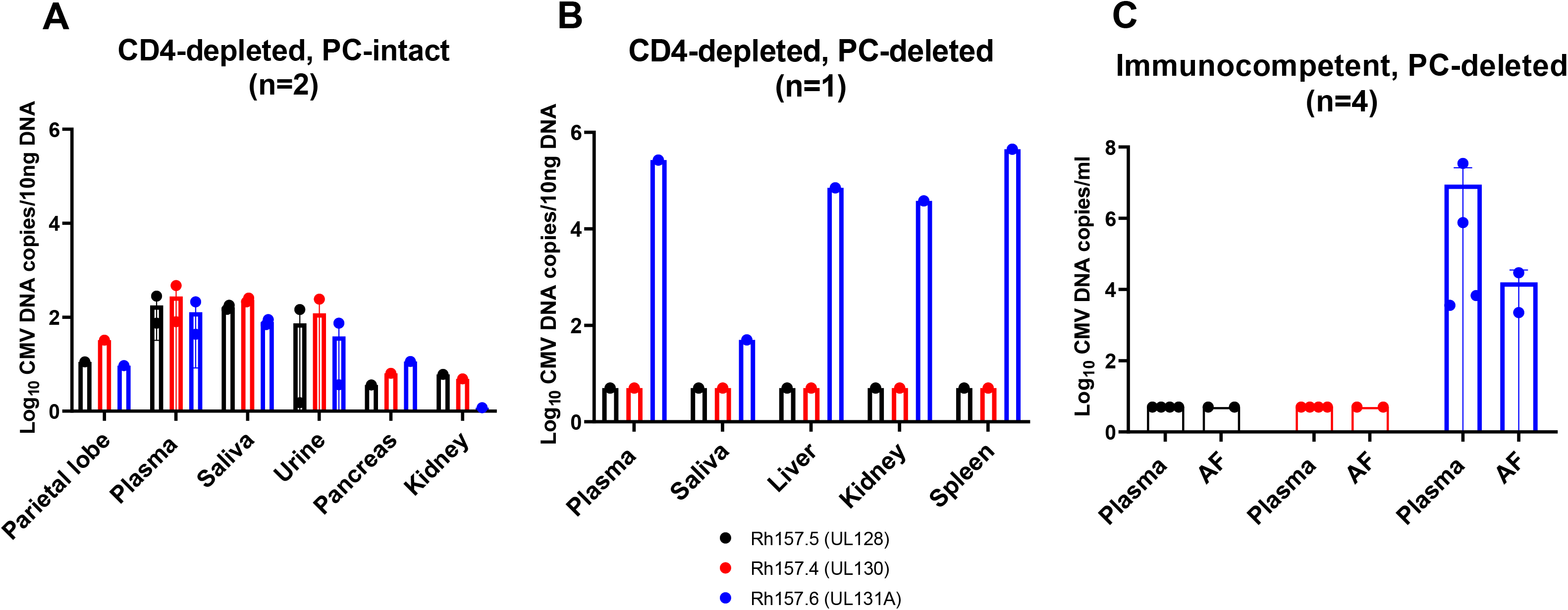
No Rh157.5 (UL128) and Rh157.4 (UL130) detection in RMs inoculated with PC-deleted RhCMV. PC subunit Rh157.5 (UL128), Rh157.4 (UL130), and Rh157.6 (UL131A) gene expression by QPCR. PC subunit gene expression of CD4-depleted dams inoculated with PC-intact RhCMV (n=2) (A), CD4-depleted dams inoculated with PC-deleted RhCMV (n=1) (B), and immunocompetent dams inoculated with PC-deleted RhCMV (n=4) (C). Maternal fluid or tissue samples high RhCMV DNA copies and enough quantity were selected to run Rh157.5/Rh157.4/Rh157.6 QPCR. Only 2 immunocompetent dams inoculated with PC-deleted RhCMV had virus detection in amniotic fluid, so QPCR was performed on these 2 amniotic fluid samples. Data shown as the mean RhCMV copy number of 3 individual replicates. Each dot indicate each individual animal at selected timepoint. Black: Rh157.5 (UL128) gene; red: Rh157.4 (UL130) gene; blue: Rh157.6 (UL131A) gene.

**Sup Figure 4.**
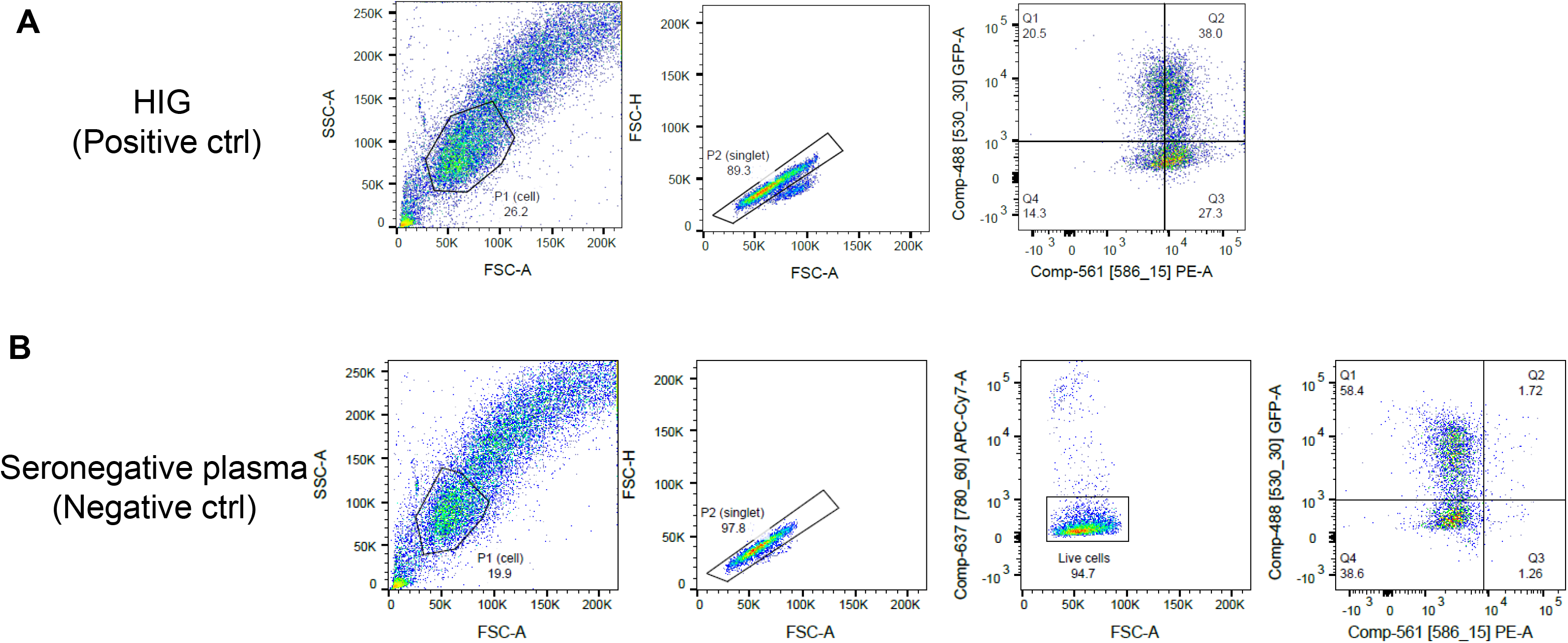
Gating strategy of gH-transfected cell binding assay. Gating strategy of gH-transfected cell binding assay for the positive control, rhesus hyperimmunoglobulin (A), and negative control, seronegative rhesus plasma (B). Singlet 293T cell population was gated by FSC-A and SSC-A, and FSC-A and FSC-H, accordingly. This population was next gated for live cells (APC-Cy7-population). Live cells were later gated based on GFP and PE expression in quadruples. The rhesus plasma IgG binding to cell-associated gH was measured by GFP+PE+ live cell population.

**Sup Figure 5.**
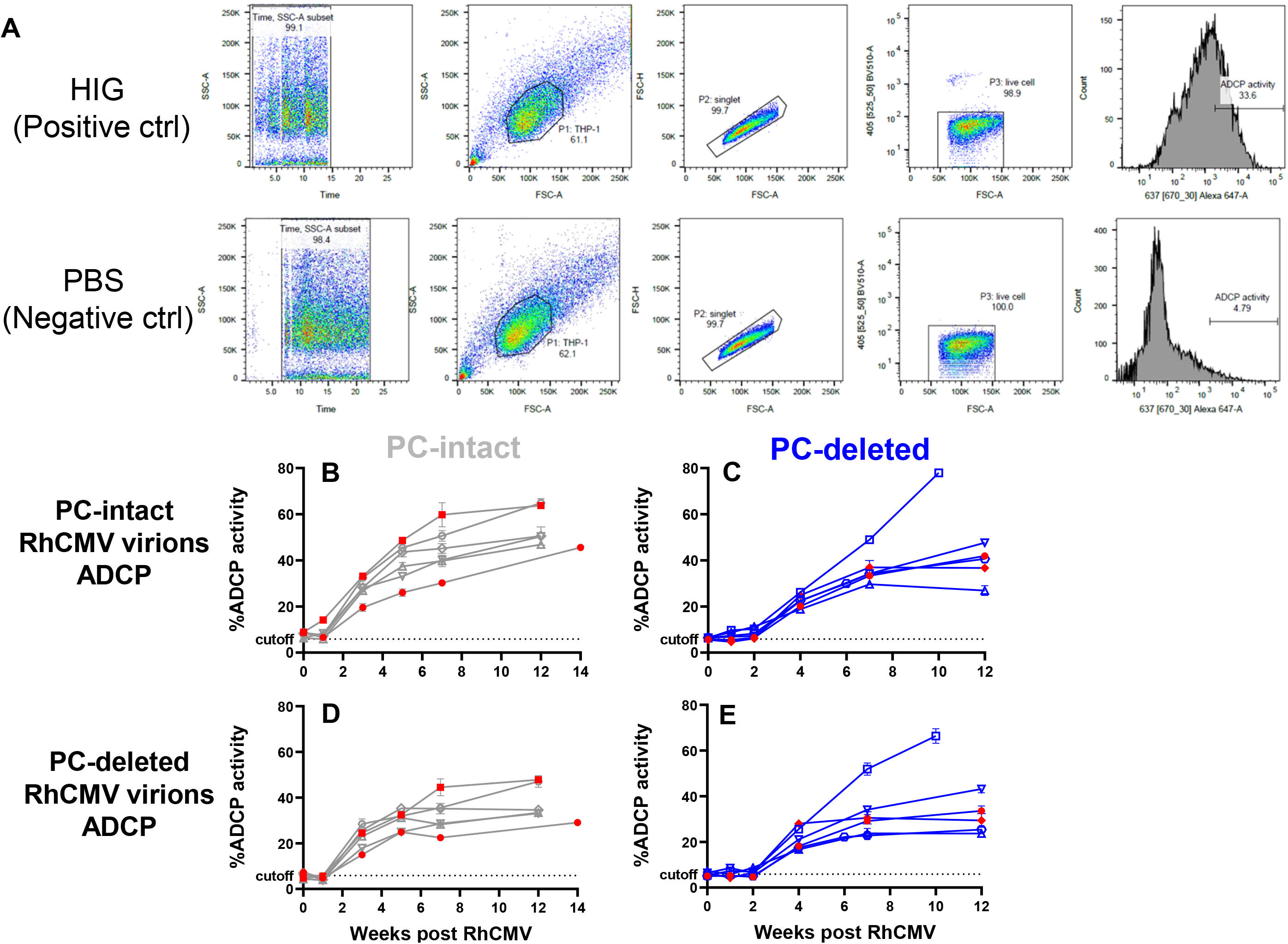
Gating strategy of antibody-dependent phagocytosis (ADCP) response and kinetics across all trimesters in immunocompetent dams. (A) Gating strategy of ADCP response for positive control (rhesus hyperimmunoglobulin) and negative control (PBS). Time gate was applied to select cells. Singlet 293T cell population was gated by FSC-A and SSC-A, and FSC-A and FSC-H, accordingly. This population was next gated for live cells (BV505-population). ADCP activity was determined by PE expression on live cells. PBS control was set to be <5% to determine PE+ population. (B-E) ADCP activity against PC-intact RhCMV whole virions (B,C) or PC-deleted RhCMV whole virions (D, E). Data points are shown as the %ADCP activity of each animal at selected timepoints. The dotted line indicates the assay cutoff, determined by (%ADCP activity of pooled seronegative rhesus plasma) mean+3 S.D. Gray open icons with gray line: dams inoculated with PC-intact RhCMV, no virus detected in AF; red closed icons with gray line: dams inoculated with PC-intact RhCMV, virus detected in AF; Blue open icons with blue line: dams inoculated with PC-deleted RhCMV, no virus detected in AF; red closed icons with blue line: dams inoculated with PC-deleted RhCMV, virus detected in AF.

**Sup Figure 6.**
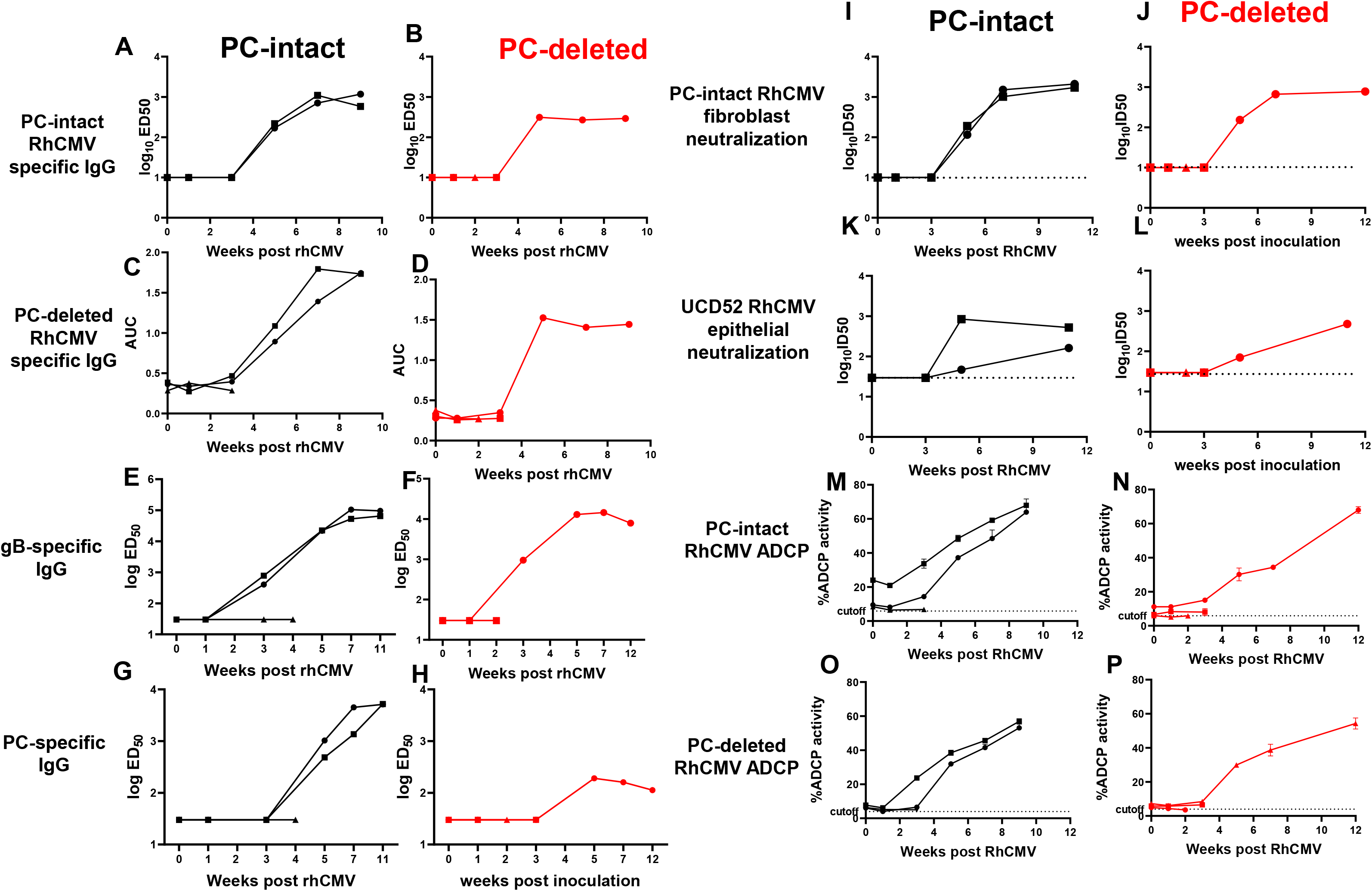
IgG binding, neutralizing antibody titers, and antibody-dependent phagocytosis response in CD4-depleted dams. (A-H) Rhesus plasma IgG binding kinetics to RhCMV whole virions and glycoproteins post virus inoculation. Rhesus plasma IgG binding to PC-intact RhCMV whole virions (A,B), PC-deleted RhCMV whole virions (C,D), RhCMV gB (E-F), and RhCMV PC (G-H) by ELISA. Data points are shown as the ED50 of each animal at selected timepoints. (I-L) Rhesus plasma IgG neutralization on fibroblasts against PC-intact RhCMV (I-J) and on epithelial cells against UCD52 strain (K-L). Data points are shown as the ID50 of each animal at selected timepoints. The dotted line indicates the assay limit of detection. (M-P) Rhesus plasma IgG % ADCP activity against PC-intact RhCMV whole virions (M-N) and PC-deleted RhCMV whole virions (O-P). Data points are shown as the %ADCP activity of each animal at selected timepoints by flow cytometry. Black closed icons: CD4-depleted dams inoculated with PC-intact RhCMV; red closed icons: CD4-depleted dams inoculated with PC-deleted RhCMV.

**Sup Figure 7.**
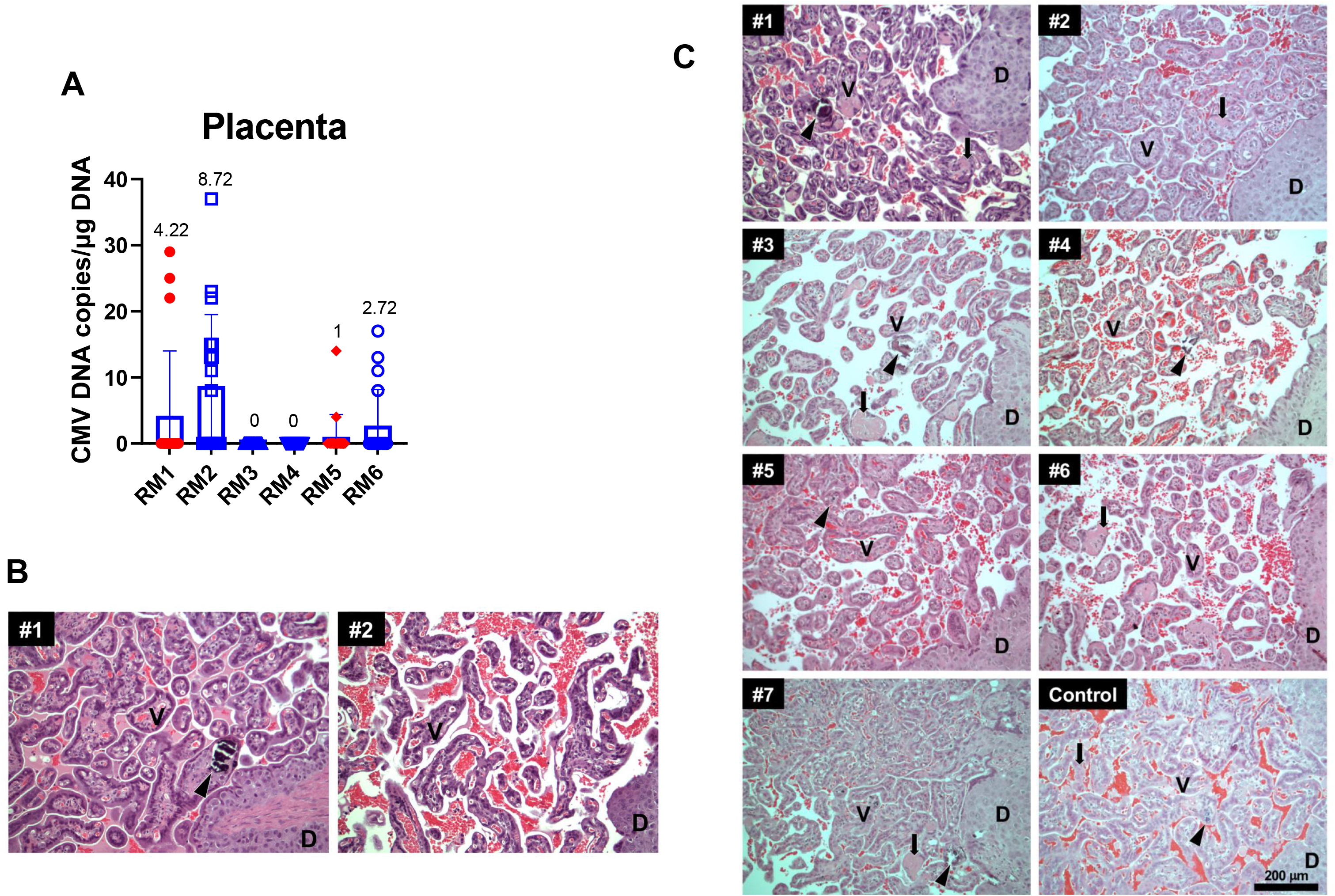
Placenta RhCMVDNA copies and histomorphology of immunocompetent dams inoculated with PC-deleted RhCMV. (A) RhCMV DNA copy numbers in placenta, collected from 6 immunocompetent dams inoculated with PC-deleted RhCMV. Data shown as the mean RhCMV copy number of 18 individual replicates. Blue open icons: dams inoculated with PC-deleted RhCMV, no virus detected in AF; red closed icons: dams inoculated with PC-intact RhCMV, virus detected in AF. (B-C) Histomorphology of placentas. Sections of placenta stained with hematoxylin and eosin (H&E) are shown from two gravid RMs that were CD4-depleted one week prior to administration of PC-intact RhCMV (#1, #2) (B). Sections are also shown from pregnant RMs inoculated with PC-deleted RhCMV (#1, CD4-depleted; #2 to #7, immunocompetent), compared to untreated controls (one example shown) (C). There were no significant findings in the placentas of animals inoculated with RhCMV. All RhCMV and control placentas showed mature normovascular chorionic villi (V), some with scattered calcifications (arrowheads) and fibrin deposition (arrows). D=decidua.

